# Catalyst: Fast and flexible modeling of reaction networks

**DOI:** 10.1101/2022.07.30.502135

**Authors:** Torkel E. Loman, Yingbo Ma, Vasily Ilin, Shashi Gowda, Niklas Korsbo, Nikhil Yewale, Chris Rackauckas, Samuel A. Isaacson

## Abstract

We introduce Catalyst.jl, a flexible and feature-filled Julia library for modeling and high-performance simulation of chemical reaction networks (CRNs). Catalyst supports simulating stochastic chemical kinetics (jump process), chemical Langevin equation (stochastic differential equation), and reaction rate equation (ordinary differential equation) representations for CRNs. Through comprehensive benchmarks, we demonstrate that Catalyst simulation runtimes are often one to two orders of magnitude faster than other popular tools. More broadly, Catalyst acts as both a domain-specific language and an intermediate representation for symbolically encoding CRN models as Julia-native objects. This enables a pipeline of symbolically specifying, analyzing, and modifying CRNs; converting Catalyst models to symbolic representations of concrete mathematical models; and generating compiled code for numerical solvers. Leveraging ModelingToolkit.jl and Symbolics.jl, Catalyst models can be analyzed, simplified, and compiled into optimized representations for use in numerical solvers. Finally, we demonstrate Catalyst’s broad extensibility and composability by highlighting how it can compose with a variety of Julia libraries, and how existing open-source biological modeling projects have extended its intermediate representation.

## 1 Introduction

Chemical reaction network (CRN) models are used across a variety of fields, including the biological sciences, epidemiology, physical chemistry, combustion modeling, and pharmacology [1–7]. At their core, they combine a set of species (defining a system’s state) with a set of reaction events (rates for reactions occurring combined with rules for altering the system’s state when a reaction occurs). One advantage of formulating a biological model as a CRN is that these can be simulated according to several well-defined mathematical representations, representing different physical scales at which reaction processes can be studied. For example, the *reaction rate equation* (RRE) is a macroscopic system of ordinary differential equations (ODEs), providing a deterministic model of chemical reaction processes. Similarly, the *chemical Langevin equation* (CLE) is a system of stochastic differential equations (SDEs), providing a more microscopic model that can capture certain types of fluctuations in reaction processes [8]. Finally, stochastic chemical kinetics, typically simulated with the *Gillespie algorithm* (as well as modifications to, and improvements of, it), provides an even more microscopic model, that captures both stochasticity and discreteness of populations in chemical reaction processes [9,10]. That a CRN can be unambiguously represented using these models forms the basis of several CRN modeling tools [11–26]. Here we present a new modeling tool for CRNs, Catalyst.jl, which we believe offers a unique set of advantages for both inexperienced and experienced modelers.

Catalyst’s defining trait, which sets it apart from other popular CRN modeling packages, is that it represents models in an entirely symbolic manner, accessible via standard Julia language programs. This permits algebraic manipulation and simplification of the models, either by the user, or by other tools. Once a CRN has been defined, it is stored in a symbolic *intermediate representation* (IR). This IR is the target of methods that provide functionality to Catalyst, including numerical solvers for both continuous ODEs and SDEs, as well as discrete Gillespie-style stochastic simulation algorithms (SSAs). As Catalyst’s symbolic representations can be converted to compiled Julia functions, it can be easily used with a variety of Julia libraries. These include packages for parameter fitting, sensitivity analysis, steady state finding, and bifurcation analysis. To simplify model implementation, Catalyst provides a *domain-specific language* (DSL) that allows users to declare CRN models using classic chemical reaction notation. Finally, Catalyst also provides a comprehensive API to enable programmatic manipulation and combination of models, combined with functionality for analyzing and simplifying CRNs (such as detection of conservation laws and elimination of conserved species).

Catalyst is implemented in Julia, a relatively recent (version 1.0 released in August 2018) open-source programming language for scientific computing. Its combination of high performance and user-friendliness makes it highly promising [27,28]. Julia has grown quickly, with a highly developed ecosystem of packages for scientific simulation. This includes the many packages provide by the Scientific Machine Learning (SciML) organization, of which Catalyst is a part. SciML, through its ModelingToolkit.jl package, provides the IR on which Catalyst is based [29]. This IR is used across the organization’s projects, providing a target structure both for model-generation tools (such as Catalyst), and tools that provide additional functionality. ModelingToolkit symbolic models leverage the Symbolics.jl [30] *computer algebraic system* (CAS), enabling them to be represented in a symbolic manner. Simulations of ModelingToolkit-based models are typically carried out using DifferentialEquations.jl, perhaps the largest software package of state-of-the-art, high-performance numerical solvers for ODEs, SDEs, and jump processes [31].

Several existing modeling packages provide overlapping functionality with Catalyst. COPASI is a well known and popular software that enables both deterministic and stochastic CRN modeling, as well as many auxiliary features (such as parameter fitting and sensitivity analysis) [12]. BioNetGen is another such software suite, currently available as a Visual Studio Code extension, that is built around the popular BioNetGen language for easily specifying complex reaction network models [21]. It provides options for model creation, network simulation, and network free-modeling. Another popular tool, VCell, provides extensive functionality, via an intuitive graphical interface [11]. Finally, Tellurium ties together a range of tools to be used in a Python environment, allowing CRN models to be created using the Antimony DSL, and simulated using the libRoadRunner numeric solver suite [15,23,24,32]. Other modeling softwares include GINsim, CellNOpt, GillesPy2, and Matlab’s SimBiology toolbox [13,16,33].

Several of these packages are primarily designed around a GUI-based workflow (BioNetGen, COPASI, and VCell). In constrast, Catalyst is DSL and API-based, with simulation and analysis of models carried out via Julia scripts. A typical Catalyst workflow therefore requires users to write Julia language scripts instead of using a GUI-based interface, but also enables users to easily integrate Catalyst models with a large variety of other Julia libraries. Catalyst also has immediate access to a more extensive set of numerical solvers for ODEs, SDEs, and SSAs. In this paper, we demonstrate that using these solvers, Catalyst’s simulation speed often outperforms the other tools by more than one order of magnitude. Catalyst has the ability to include Julia-native functions within rate laws and stoichiometric expressions, and to include coupled ODE or algebraic constraint equations for reaction rate equation models (potentially resulting in *differential-algebraic equation* (DAE) models). For example, to encode bursty reactions stoichiometric coefficients can be defined using standard Julia functions that sample from a random variable distribution. Similarly, rate-laws can include data-driven modeling terms (e.g. neural networks) constructed via Julia libraries such as Surrogates.jl, SciMLSensitivity.jl, and DiffEqFlux.jl. Moreover, Catalyst generates differentiable models, which can be easily incorporated into higher-level Julia codes that require automatic differentiation [34] and composed with other Julia libraries. One current limitation of Catalyst is that in contrast to BioNetGen, COPASI, and GillesPy2, Catalyst can not generate inputs for hybrid and *τ* -leaping solvers, though adding support for these features is planned.

In the next sections we overview a basic workflow for using Catalyst to define and simulate CRNs; overview how Catalyst performs relative to several popular CRN modeling packages for solving ODEs and simulating stochastic chemical kinetics models; discuss Catalyst’s symbolic representation of CRNs, Catalyst’s network analysis functionality, and how it can compose with other Julia packages; and introduce some of the higher-level applications in which Catalyst models can be easily embedded.

## 2 Results

### 2.1 The Catalyst DSL enables models to be created using chemical reaction notation

Catalyst offers several ways to define a CRN model, with the most effortless option being the @reaction network DSL. This feature extends the natural Julia syntax via a macro, allowing users to declare CRN models using classic chemical reaction notation (as opposed to declaring models using equations, or by declaring reactions implicitly or through functions). This alternative notation makes scripts more human-legible, and greatly reduces code length (simplifying both script writing and debugging). Using the DSL, the CRN’s chemical reactions are listed, each preceded by its reaction rate (Fig. 1). From this, the system’s species and parameters are automatically extracted and a `ReactionSystem` IR structure is created (which can be used as input to e.g. numerical simulators).

**Figure 1:**
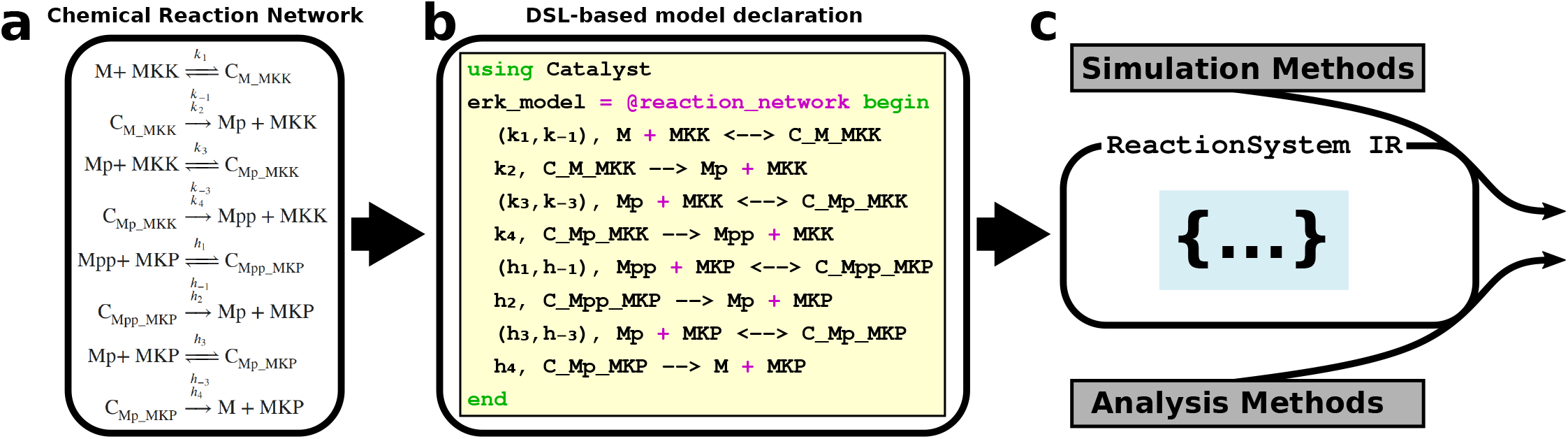
Catalyst connects an intuitive domain-specific language with a well-supported intermediate representation. The extracellular signal-regulated kinase (ERK) network is important to the regulation of many cellular functions, and its disruption has been implicated in cancer [35]. (a) A CRN representation of the ERK network. (b) A model of the ERK CRN can be implemented in Julia through the Catalyst DSL, using code very similar to the actual CRN representation. (c) From this code, the DSL generates a `ReactionSystem` intermediate representation (IR) that is the target structure for a range of supported simulation and analysis methods.

To facilitate a more concise notation, similar reactions (e.g. several degradation events) can be bundled together. Each reaction rate can either be a constant, a parameter, or a function. Predefined Michaelis–Menten and Hill functions are provided by Catalyst, but any user-defined Julia function can be used to define a rate. Both parametric and non-integer stoichiometric coefficients are possible. There are also several non-DSL methods for model creation. They include loading networks from files via SBMLToolkit.jl [36] (for SBML files) and ReactionNetworkImporters.jl [37] (for BioNetGen generated.net files). CRNs can also be created via defining symbolic variables via the combined Catalyst/ModelingToolkit API, and directly building `ReactionSystem`s from collections of `Reaction` structures. This enables programmatic definition of CRNs, making it possible to create large models by iterating through a relatively small number of rules within standard Julia scripts.

### 2.2 Catalyst models can be simulated using a wide range of high-performance methods

Numerical simulations of Catalyst models are generally carried out using the DifferentialEquations.jl package [31]. It contains a large number of numerical solvers and a wide range of additional features (such as event handling, support for GPUs and threading, flexibility in choice of linear solvers for stiff integrators, and more). The package is highly competitive, often outperforming packages written in C and Fortran [31]. Simulation syntax is straightforward, and output solutions can be plotted using the Plots.jl package [41] via a recipe that allows users to easily select the species and times to display. CRNs can be translated and simulated using the ODE-based RREs, the SDE-based CLE, and through discrete SSAs (Fig. 2).

**Figure 2:**
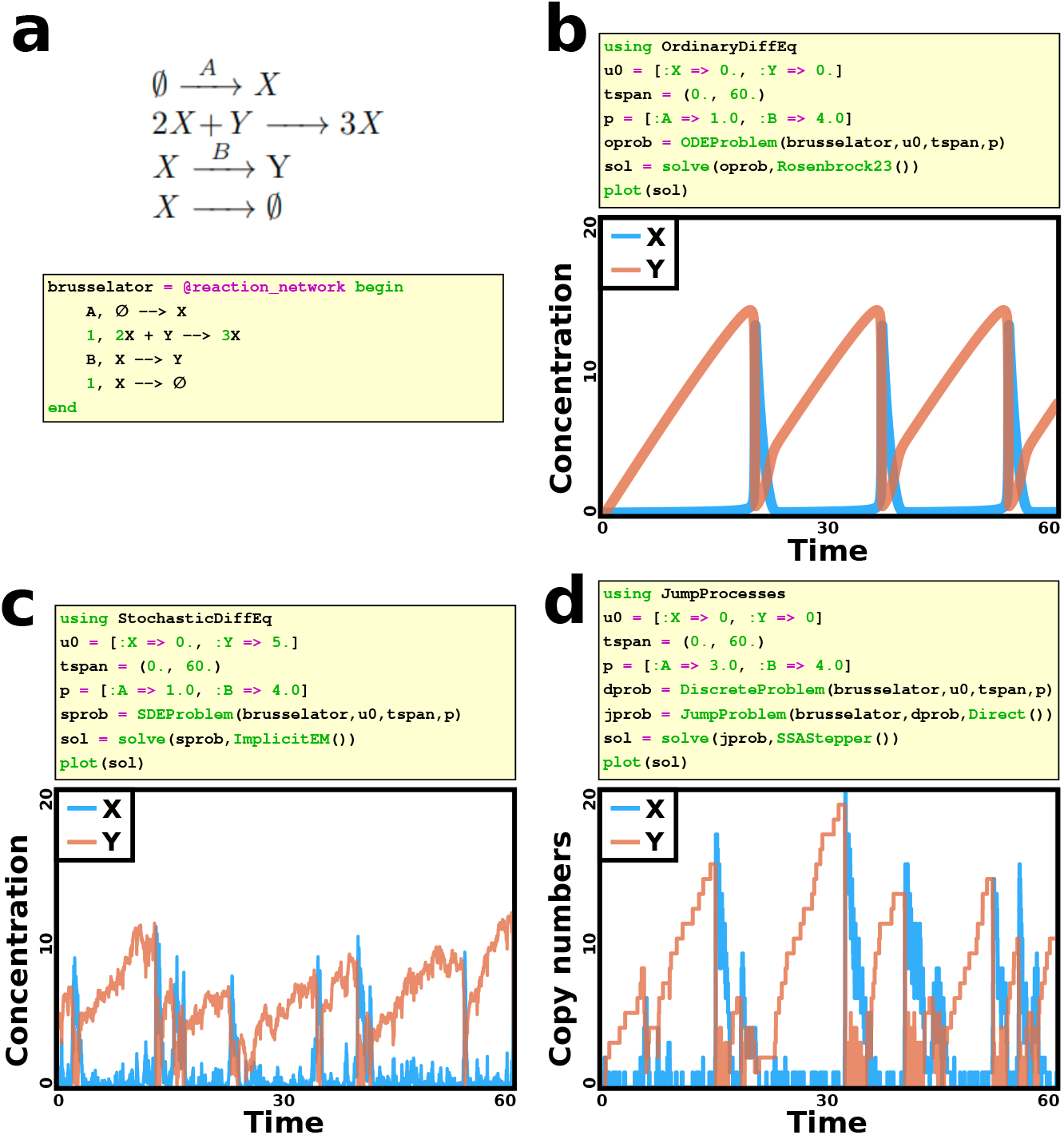
Catalyst models can be simulated using both deterministic and stochastic interpretations. (a) The Brusselator network contains two species (X and Y) and two parameters (A and B, in practical implementation these are species present in excess, but they can in practice be considered parameters) [38,39]. Here, we show the four reactions of the Brusselator CRN, and its implementation using the Catalyst DSL. (b-d) Simulations of models for the Brusselator at the three physical scales supported by Catalyst (RRE, CLE, SSA). Post-processing has been carried out on the plots to improve their visualization in this article’s format. (b) While *B >* 1 + *A*^2^, the deterministic model exhibits a limit cycle. This is confirmed using ODE RRE simulations. (c) The model can also be simulated using the stochastic CLE interpretation. (d) Finally, the discrete, stochastic, jump process interpretation is simulated via Gillespie’s direct method. The system displays a limit cycle even though *B <* 1 + *A*^2^, confirming the well known phenomenon of noise induced oscillations [40].

To demonstrate the performance of these solvers, we benchmarked simulations of CRN models using a range of CRN modeling tools (BioNetGen, Catalyst, COPASI, GillesPy2, and Matlab’s SimBiology toolbox). These tools were selected as they are popular and highly cited, well documented, scriptable for running benchmark studies, and actively maintained. The Matlab SimBiology toolbox was selected due to the enduring popularity of the Matlab language. Overall, they provide a representative sample of the broader chemical reaction network modeling software ecosystem. We used both ODE simulations and discrete SSAs. Fewer packages permit SDE simulations, hence such simulations were not benchmarked. We note, however, that DifferentialEquations’ SDE solvers are highly performative [48]. When comparing a range of models, from small to large, we see that Catalyst typically outperforms the other packages, often by at least an order of magnitude (Fig. 3). For the ODE benchmarks, to try to provide as fair a comparison as possible, identical absolute and relative tolerances were used for all simulations. Furthermore, in Supplementary Fig. E.1 we demonstrate the relation between simulation time and actual error across the Julia solvers, lsoda, and CVODE (with the native Julia solvers typically having smaller errors as compared to lsoda and CVODE for any given tolerance). All SSA methods tested generate exact realizations, in the sense that they should each give statistics consistent with the underlying Chemical Master Equation of the model [49], and their simulation times are hence directly comparable. Here, the wide range of methods provided by the JumpProcesses.jl package [50], a component of DifferentialEquations, enables Catalyst to outperform the other packages (most of which only provide Gillespie’s direct method or its sorting direct variant [51]).

**Figure 3:**
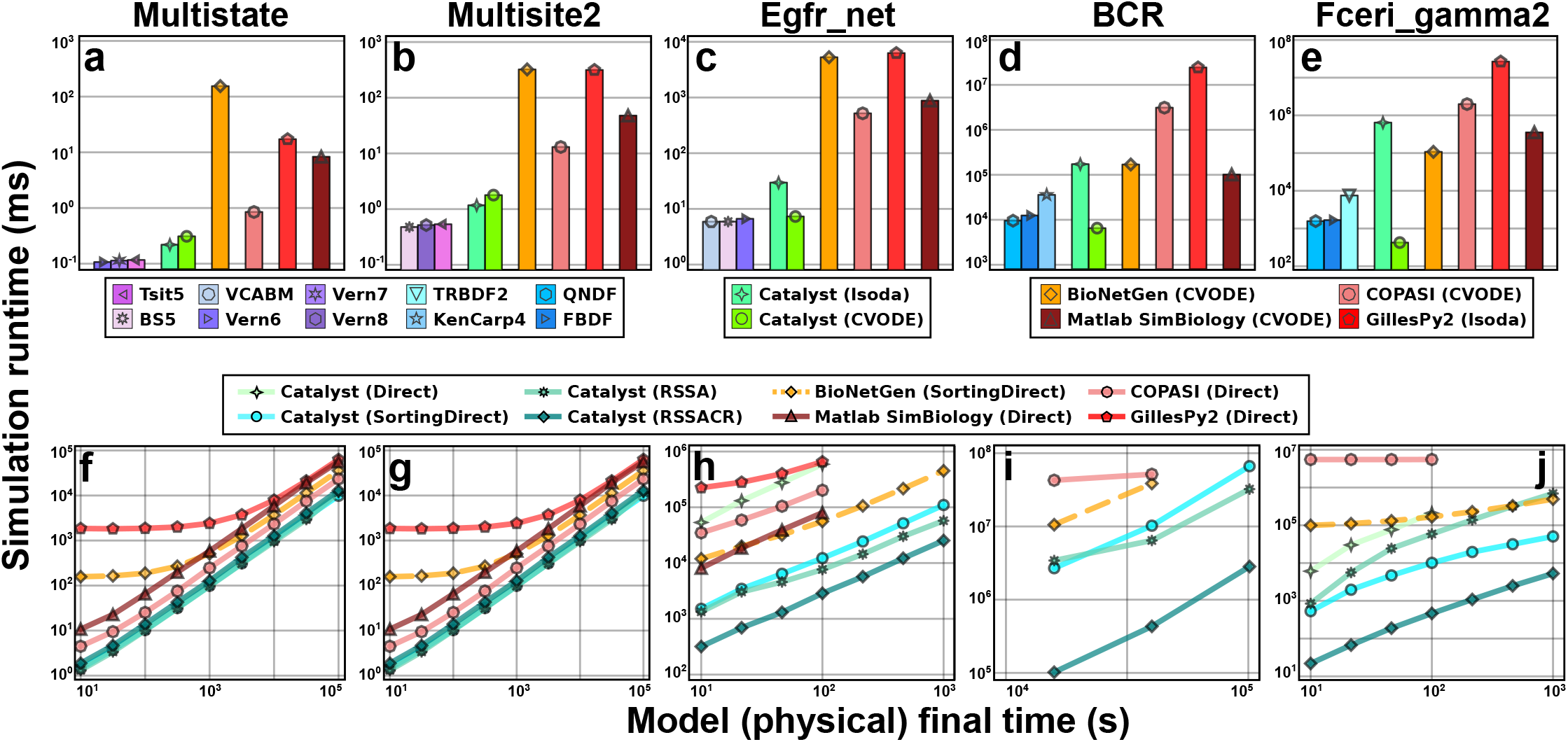
Simulations of Catalyst models outperform those of other modeling packages. Benchmarks of simulation runtimes for Catalyst and four other modeling packages (BioNetGen, COPASI, GillesPy2, and Matlab SimBiology). The benchmarks were run on the multi-state (Multistate, 9 species and 18 reactions [42]), multi-site (Multisite 2, 66 species and 288 reactions [43]), epidermal growth factor receptor signalling (Egfr_net, 356 species and 3749 reactions [44]), B-cell receptor (1122 species and 24388 reactions [45]), and high-affinity human IgE receptor signalling (Fceri gamma2, 3744 species and 58276 reactions [46]) models. (a-e) Benchmarks of deterministic RRE ODE simulations of the five models. Each bar shows, for a given method, the runtime to simulate the model (to steady-state for those that approach a steady-state). For Catalyst, we show the three best-performing native Julia methods, as well as the performance of lsoda and CVODE. For each of the other tools, we show its best-performing method. Identical values for absolute and relative tolerance are used across all packages and methods. For each benchmark, the method options used can be found in table 3, the exact benchmark times in Table 4, and further details on the solver options for each tool in Supplementary Section B. While this figure only contains the most performant methods, a full list of methods investigated can be found in Supplementary Section B, with their results described in Supplementary Figs. D.1 and D.2. (f-j) Benchmarks of stochastic chemical kinetics SSA simulations of the five models. Via JumpProcesses.jl, Catalyst can use several different algorithms (e.g. Direct, Sorting Direct, RSSA, and RSSACR above) for exact Gillespie simulations. Here, the simulation runtime is plotted against the (physical) final time of the simulation. Due to their long runtimes, some tools were not benchmarked for the largest models. We note that, in [47], it was remarked that BioNetGen (dashed orange lines) use a pseudo-random number generator in SSAs that, while fast, is of lower quality than many (slower) modern generators such as Mersenne Twister. For full details on benchmarks, see Section 4.1.

In contrast to the exact SSA methods, timestep-based ODE integrators typically provide a variety of numerical parameters, such as error tolerances and configuration options for implicit solvers (i.e. how to calculate Jacobians, how to solve linear and nonlinear systems, etc). ODE simulation performance then depends on which combinations of options are used with a given solver. Here, we limit ourselves to trying combinations of numeric solvers (Julia-native solvers for comparing performance of Catalyst-generated models, and lsoda and/or CVODE for comparisons between tools), methods for Jacobian computation and representation (automatic differentiation, finite differences, or symbolic computation, and dense vs. sparse representations), linear solvers (LU, GMRES, or KLU), and whether to use a preconditioner or not when using GMRES. The non-Catalyst simulators generally provide limited ability to change these options, in which case only the default was used in benchmarking. In contrast, the DifferentialEquations.jl solvers that Catalyst utilise, while they do not require the user to set these options, do give them full control to do so. Full documentation is available at [52]. The details of the most performant options we used for each tool and model are provided in Table 3. A list of all benchmarks we carried out (for various combinations of tool, method, and options) is provided in Supplementary Section B, with their results described in Supplementary Figs. D.1 and D.2. Finally, the benchmarking process is described in more detail in Section 4.1.

The observed performance results for Catalyst-generated models arise from a variety of factors. For example, Catalyst inlines all mass action reaction terms for ODE models within a single generated function that evaluates the ODE derivative. This provides opportunities for the compiler to optimise expression evaluation, and avoids the overhead of repeatedly calling non-inlined functions to evaluate such terms. For the largest ODE models, Catalyst and ModelingToolkit’s support for generating explicit sparse Jacobians led to significant performance improvements when using the CVODE solver, see Section 4.1 and the supplement. For jump process SSA simulations, Catalyst uses a sparse reaction specification that automatically analyses each reaction, and then classifies the reaction into the most performant but physically valid representation supported by JumpProcesses.jl (corresponding to jumps with general time-varying intensities, jumps with general rate expressions but for which the intensity is constant between the occurrence of two jumps, and jumps for which the intensity is a mass action type rate law). This enables JumpProcesses.jl to avoid the overhead of calling a large collection of user-provided functions via pointers, by using a single pre-defined and inlined function to evaluate individual mass action reaction intensities, while still supporting calling general user rate functions via pointers (for non-mass action rate laws). These Catalyst-specific features, when coupled to the large variety of solvers in DifferentialEquations.jl and broad flexibility in tuning solver components (i.e. different Jacobian and jump representations, flexibility in choice of linear solvers, etc.), help enable Catalyst’s observed performance.

### 2.3 Catalyst enables composable, symbolic modeling of CRNs

Catalyst’s primary feature is that its models are represented using a CAS, enabling them to be algebraically manipulated. Examples of how this is utilised include automatic computation of system Jacobians, calculation and elimination of conservation laws, and simplification of generated symbolic DAE models via ModelingToolkit’s symbolic analysis tooling. These techniques can help speed up numeric simulations, while also facilitating higher level analysis. One example is enabling users to generate ODE models with non-singular Jacobians via the elimination of conservation laws, which can aid steady-state analysis tooling. Catalyst also provides a network analysis API, enabling the calculation of a variety of network properties beyond conservation laws, including linkage classes, weak reversibility, and deficiency indices.

Catalyst’s symbolic representation permits model internals to be freely extracted, investigated, and manipulated, giving the user full control over their models (Fig. 4). This enables various forms of programmatic model creation, extension and composition. Model structures that occur repetitively can be duplicated, and disjoint models can be connected together. For example, such functionality can be used to model a population of cells, each with defined neighbours, where each cell can be assigned a duplicate of the same simple CRN. The CRNs within each cell can then be connected to those of its neighbours, enabling models with spatial structures. Similarly, one could define a collection of genetic modules, and then compose such modules together into a larger gene regulatory network.

**Figure 4:**
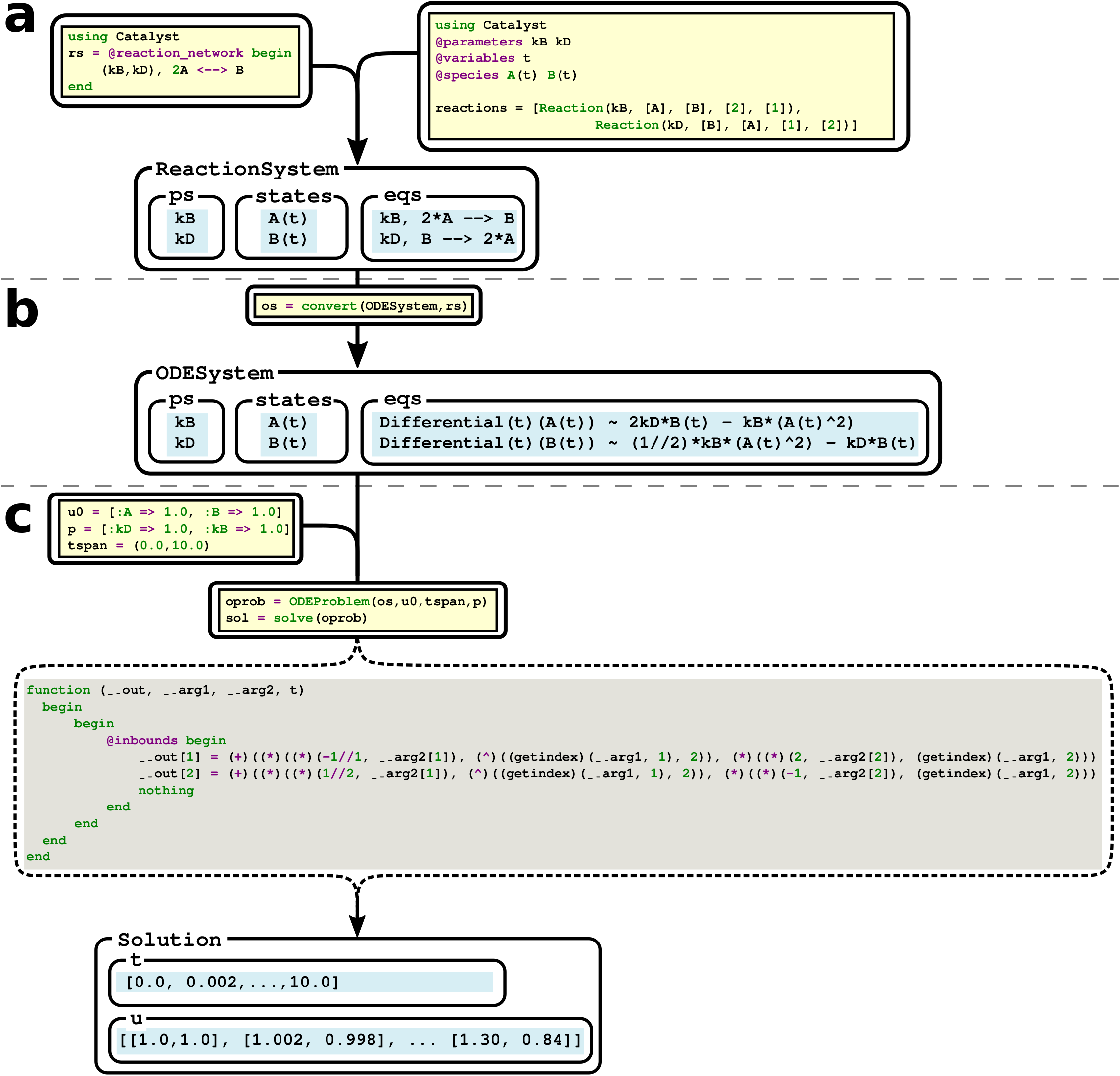
The simulation pipeline of a Catalyst model, with internal intermediates displayed. Code as written by the user (yellow background), and as generated internally by Catalyst and ModelingToolkit (blue and grey backgrounds respectively) are shown, in addition to the generated structures and their fields (blue background, some of the internal fields are omitted in all displayed structures). (a) A symbolic `ReactionSystem` for a reversible dimerisation reaction is created using either the DSL, or programmatically using the Symbolics computer algebraic system. (b) The `ReactionSystem` can be converted into a ModelingToolkit `ODESystem` structure, corresponding to a symbolic RRE ODE model. (c) By providing initial conditions, parameter values, and a time span, the `ODESystem` can be simulated, generating an output solution. The generated (internal) Julia code for evaluating the derivatives defining the ODEs, which gets compiled and is input to the ODE solver, is displayed in grey. At each step, the user has the ability to investigate and manipulate the generated structures.

Catalyst is highly flexible in the allowed Julia functions that can be used in defining rates, rate laws, or stoichiometry coefficients. This means that while reaction rates and rate laws are typically constants, parameters, or simple functions, e.g. Hill functions, they may also include other terms, such as neural networks or data-driven, empirically defined, Julia functions. Likewise, stoichiometric coefficients can be random variables by defining them as a symbolic variable, and setting that variable equal to a Julia function sampling the appropriate probability distribution. Such functionality can be utilized, for example, to model transcriptional bursting [53], where the produced mRNA copy-numbers are random variables. Finally, standard Catalyst-generated ODE and SDE models are differentiable, in that the generated codes can be used in higher-level packages that rely on automatic differentiation [34]. In this way Catalyst-generated models can be used in machine-learning based analyses.

That Catalyst gives full access to its model internals, combined with its composability, allows other packages to easily integrate into, and build upon, it. Indeed, this is already being utilised by independent package developers. The Moment Closure.jl Julia package, which implements several techniques for moment closure approximations, is built to be deployed on Catalyst models [54]. It can generate symbolic finite-dimensional ODE system approximations to the full, infinite system of moment equations associated with the chemical master equation. These symbolic approximations can then be compiled and solved via ModelingToolkit in a similar manner to how Catalyst’s generated RRE ODE models are handled. Similarly, FiniteStateProjection.jl [55] builds upon Catalyst and ModelingToolkit to enable the numerical solution of the chemical master equation, while DelaySSAToolkit.jl [56] can accept Catalyst models as input to its SSAs that handle stochastic chemical kinetics models with delays. Another example of how Catalyst’s flexibility enables its integration into the Julia ecosystem is that CRNs with polynomial ODEs (a condition that holds for pure mass action systems) can be easily converted to symbolic steady-state systems of polynomial equations. This enables polynomial methods, such as homotopy continuation, to be employed on Catalyst models. Here, homotopy continuation (implemented by the HomotopyContinuation.jl Julia package) can be used to reliably compute all roots of a polynomial system [57]. This is an effective approach for finding multiple steady states of a system. When the CRN contains Hill functions (with integer exponents), by multiplying by the denominators, one generates a polynomial system with identical roots to the original, on which homotopy continuation can still be used.

### 2.4 Catalyst models are compatible with a wide range of ancillary tools and methods

The Julia SciML, and broader Julia, ecosystem offers a wide range of techniques for working with models and data based around the IR that Catalyst produces (Fig. 5). While the reactions that constitute a CRN are often known in developing a model, system parameters (these typically correspond to reaction rates) rarely are. A first step in analyzing a model is identifiability analysis, where we determine whether the parameters can be uniquely identified from the data [58]. This is enabled through the StructuralIdentifiability.jl package. In the next step, parameters can be fitted to data. This can be done using DiffEqParamEstim.jl, which provides simple functions that are easy to use. Alternatively, more powerful packages, like Optimization.jl and the Turing.jl Julia library for Bayesian analysis, offer increased flexibility for experienced users [59]. Furthermore, unknown CRN structures (such as a species’s production rate) can be approximated using neural networks and then fitted to data. This functionality is enabled by the SciMLSensitivity package [60]. More broadly, system steady states can be computed using the NLSolve.jl or HomotopyContinuation.jl Julia packages [57]; bifurcation structures can be calculated, and bifurcation diagrams generated, with the BifurcationKit.jl library [61]; and SciMLSensitivity.jl and GlobalSensitivity.jl can be used to investigate the sensitivity and uncertainty of model solutions with regard to parameters [62]. Finally, options for displaying CRNs, either as network graphs (via Graphviz) or Latex formatted equations (via Latexify.jl), also exist.

**Figure 5:**
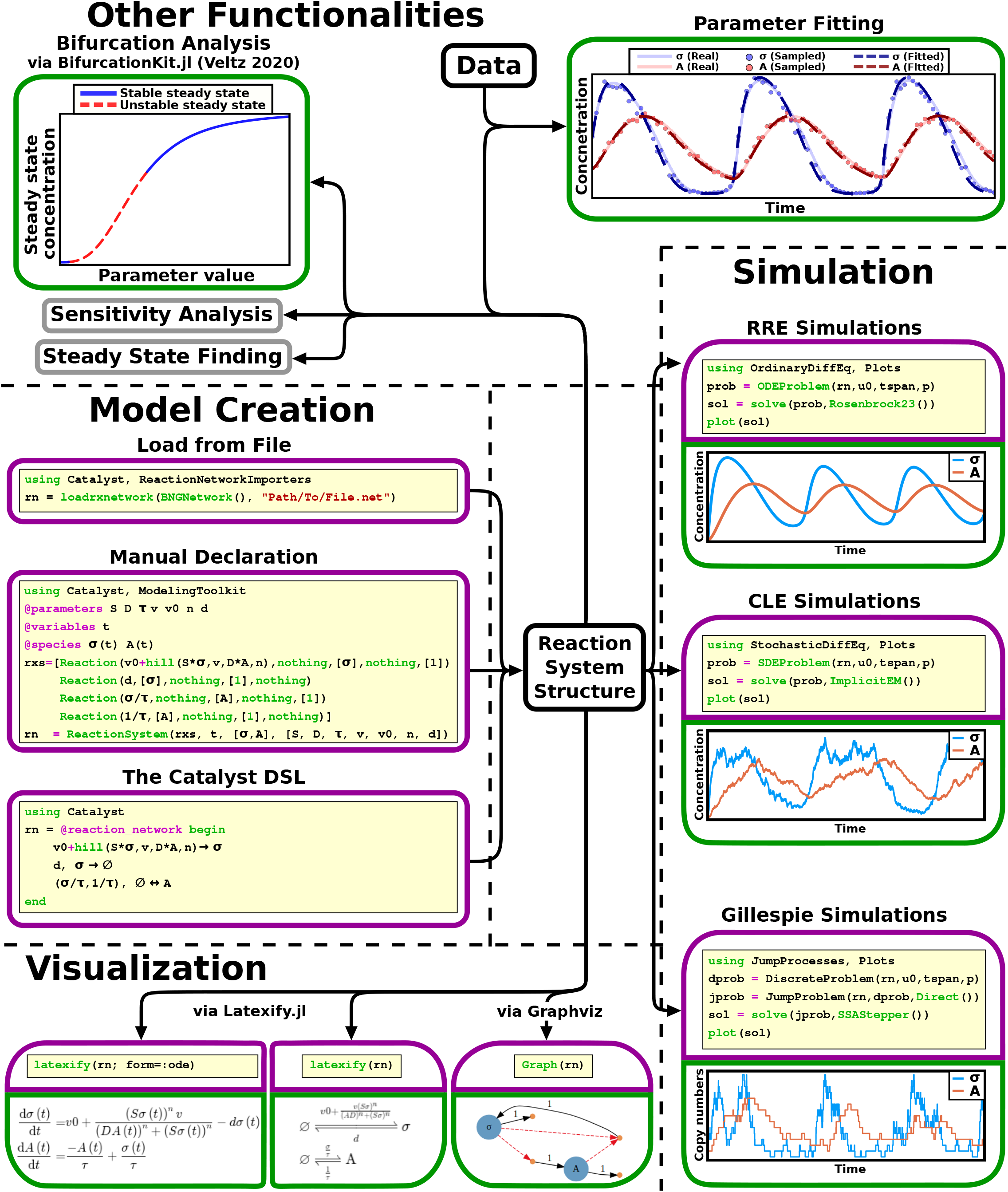
A wide range of features are available for Catalyst model analysis. A CRN model can be created either through the DSL, by manually declaring the reaction events, or by loading it from a file. The model is stored in the `ReactionSystem` IR, which can be used as input to a wide range of methods. Purple boxes indicate code written by the user, and green boxes the corresponding output. For some methods, either one, or both, boxes are omitted.

## 3 Discussion

In this article, we have introduced the Catalyst library for modeling of CRNs. It represents models through the Modeling-Toolkit.jl IR, which is used across the SciML organization and Julia ecosystem libraries, and can be automatically translated into optimized inputs for numerical simulations (RRE ODE, CLE SDE, and stochastic chemical kinetics jump process models). Our benchmarks demonstrate that Catalyst often outperforms other tools by an order of magnitude or more. Moreover, it can compose with a variety of other Julia packages, including data-driven modeling tooling (parameter fitting and model inference), and other functionality (identifiable analysis, sensitivity analysis, steady state analysis, etc). The IR is based on the Symbolics.jl CAS, enabling algebraic manipulation and simplification of Catalyst models. This can both be harnessed by the user (e.g. to create models programmatically) and by software (e.g. for automated Jacobian computations). Finally, this also enables easy connection to other Julia packages for symbolic analysis, such as enabling polynomial methods (e.g homotopy continuations) to act on CRN ODEs that have a polynomial form.

In addition to the wide range of powerful tools enabled by the combination of the ModelingToolkit IR and the Symbolics CAS, Catalyst also provides a DSL that simplifies the declaration of smaller models. Of a finalized pipeline that evaluates a model with respect to a specific scientific problem, the model declaration is typically only a minor part. However, reaching a final model often requires the production and analysis of several alternative network topologies. If the barrier to create, or modify, a model can be reduced, more topologies can be explored in a shorter time. Thus, an intuitive interface can greatly simplify the model exploration portion of a research project. By providing a DSL that reads CRN models in their most natural form, Catalyst helps to facilitate model construction. In addition, this form of declaration makes code easier to debug, as well as making it easier to understand for non-experts.

While several previous tools for CRN modeling have been primarily designed around their own interface, we have instead designed Catalyst to be called from within standard Julia programs and scripts. This is advantageous, since it allows the flexibility of analysing a model with custom code, without having to save and load simulation results to and from files. Furthermore, by integrating our tool into a larger context (SciML), support for a large number of higher-order features is provided, without requiring any separate implementation within Catalyst. This strategy, with modeling software targeting an IR (here provided by ModelingToolkit) enables modelers across widely different domains to collaborate in the development and maintenance of tools. We believe this is the ideal setting for a package like Catalyst.

Development of Catalyst is still active, with several types of additional functionality planned. This includes specialised support for spatial models, including spatial SSA solvers for the reaction-diffusion master equation, and general support for reaction models with transport on graphs at both the ODE and jump process level. A longer-term goal is to enable the specification of continuous-space reaction models with transport, and interface with Julia partial differential equation libraries to seamlessly generate such spatially-discrete ODE and jump process models. Furthermore, unlike BioNetGen, COPASI, and GillesPy2, Catalyst does not currently support hybrid methods. These allow model components to be defined at different physical scales (such as resolving some reactions via ODEs and others via jump processes) [63,64]. This, as well as *τ* -leaping-based solvers [65,66], are planned for future updates. Such hybrid approaches can help to overcome the potential negativity of solutions that can arise in *τ* -leaping and CLE-based models [67]. In the CLE case, Catalyst currently wraps rate laws within the coefficients of noise terms in absolute values to avoid square roots of negative numbers, allowing SDE solvers to continue time-stepping even when solutions become negative (following the approach in [68]). We hope to also integrate alternative modelling approaches, such as the constrained CLE [67], which avoid negativity of solutions via modification of the dynamics at the positive-negative population boundary. Finally, given Catalyst’s support for units we hope to implement functionality for automatically converting between concentration and “number of” units within system specifications by allowing users to specify compartments with associated size units.

Catalyst is available for free under the permissive MIT License. The source code can be found at https://github.com/SciML/Catalyst.jl. It is also a registered package within the Julia ecosystem and can be installed from within a Julia environment using the commands using Pkg; Pkg.add(“Catalyst”). Full documentation, including tutorials and an API, can be found at https://catalyst.sciml.ai/stable/. Issues and help requests can be raised either at the Catalyst GitHub page, on the Julia discourse forum (https://discourse.julialang.org/), or at the SciML organization’s Julia language Slack channels (`#diffeq-bridged` and `#sciml-bridged`). The library is open to pull requests from anyone who wishes to contribute to its development. Users are encouraged to engage in the project.

## 4 Methods

### 4.1 Benchmarks

Benchmarks were carried out using the five CRN models used in [47]. The .bngl files provided in [47] were used as input to BioNetGen, while COPASI, GillesPy2, and Matlab used the corresponding (BioNetGen generated) .xml files. Catalyst used the corresponding (by BioNetGen generated) .net files. The exception was the BCR model, for which we used the .bngl file from [45], rather than the one from [47]. Throughout the simulations, no observable values were saved. Where options were available to reduce solution time point save frequency, and these improved performances, these were used (Supplementary Section C). BioNetGen, COPASI, and GillesPy2 simulations were performed using their corresponding Python interfaces. To ensure the correctness of the solvers, for each combination of model, tool, method, and options, ODE and SSA simulations were carried out and the results were plotted. The plots were inspected to ensure consistency across all simulations (Supplementary Figs. F.1-F.10). Runtimes were measured using `timeit` (in Python), BenchmarkTools.jl (in Julia, [69]), and `timeit` (in Matlab). For each benchmark, the median runtime over several simulations was used (the number of simulations carried out for each benchmark, over which we took the median, is described in Table 1).

**Table 1:** Number of simulations used to calculate median simulation times. For each benchmark, we performed a number of simulations, computing their median runtime. The number of such simulations depends on the tool and model (with this number given in this table). As default, we used 10, but in some cases we needed to reduce this to enable the benchmark to be completed within a reasonable time. For Julia and Matlab benchmarks, the number of simulations was automatically determined by the `timeit` tool and the BenchmarkTools.jl package, respectively.

| Model: | Multistate | Multisite2 | Egfr_net | BCR | Fceri_gamma2 |
| --- | --- | --- | --- | --- | --- |
| BioNetGen (ODE) | 10 | 10 | 10 | 10 | 10 |
| BioNetGen (SSA) | 10 | 10 | 10 | 4 | 5 |
| COPASI (ODE) | 10 | 10 | 10 | 10 | 10 |
| COPASI (SSA) | 10 | 10 | 10 | 2 | 5 |
| GillesPy2 (ODE) | 10 | 10 | 10 | 5 | 10 |
| GillesPy2 (SSA) | 10 | 10 | 10 | 2 | 5 |

For ODE benchmarks, simulation run times were measured from the initial conditions used in [47] to the time for the model to reach its (approximate) steady state (Table 2). The exception was the BCR model, which exhibited a pulsing limit cycle behaviour. For this, we simulated it over 20,000 time units, allowing it to complete three pulse events (Supplementary Fig. F.4). For ODE simulations, for all tools, the absolute tolerance was set to 10^−9^ and the relative tolerance 10^−6^. Primarily tests were carried out using the lsoda and CVODE solvers [70,71]. However, Catalyst has access to additional ODE solvers via DifferentialEquations.jl (more specifically OrdinaryDiffEq.jl). Some of these (such as QNDF and TRBDF2) are competitive with lsoda and CVODE, hence these additional solvers were also benchmarked [72,73]. All benchmarks were carried out on the MIT supercloud HPC [74]. We used its Intel Xeon Platinum 8260 units (each node has access to 192 GB RAM and contains 48 cores, of which only a single one was used). Each benchmark was carried out on a single, exclusive, node, to ensure they were not affected by the presence of other jobs. Julia, Matlab, and Python all were set to use only a single thread, ensuring multi-threading did not affect performance (e.g. Julia solvers will automatically utilise additional available threads to speed up the linear solvers of implicit simulators). Finally, work-precision diagrams were investigated to determine the relationship between simulation time and error in the native Julia solvers (Supplementary Fig. E.1). All benchmarking code is avaiable at (Code availability) under a permissive MIT license.

**Table 2:** Final (physical) time for model steady states in ODE benchmarks. For each model, we determined the time at which it had (approximately) reached a steady state. These times were used for the ODE benchmarks in Fig. 3 and Supplementary Fig. D.1. Unlike the other models, BCR exhibits a limit cycle. Here, rather than simulating until an (approximate) steady state had been reached, we simulated it for 20,000 time units (permitting it to complete 3 pulse events, Supplementary Fig. F.4).

| Model: | Multistate | Multisite2 | Egfr_net | BCR | Fceri_gamma2 |
| --- | --- | --- | --- | --- | --- |
|  | 20 s | 2 s | 10 s | 20,000 s | 150 s |

When using CVODE or implicit solvers, Catalyst permits a range of simulation options. By default, Jacobians are computed through automatic differentiation [34]. This option can either be disabled (with the Jacobian then being automatically computed through finite differences), or an option can be set to automatically compute, and use, a symbolic Jacobian from Catalyst models. Another option enables a sparse representation of the Jacobian matrix. Furthermore, the underlying linear solver for all implicit methods can be specified. We tried both the default option (which automatically selects one), but also specified either the LapackDense (using LU), GMRES, or KLU linear solvers. When the GMRES linear solver is used, a preconditioner can be set. Here we investigated both using no preconditioner, and using an incomplete LU preconditioner (described further in Supplementary Section B). Jacobians were generated using either automatic differentiation (when either the Multistate, Multisite2, or Egfr_net models were simulated using Julia solvers) or finite differences. The exception was for the KLU linear solver, for which we used a symbolically computed Jacobian. When we used either the KLU linear solver, or preconditioned GMRES, a sparse Jacobian representation was used. Generally, the non-Catalyst tools have fewer available solvers (typically depending on CVODE) and options, however, we tried those we found available. We also note that Catalyst CVODE simulations without any options specified still compare favourably to the other tools (Supplementary Fig. D.1). The methods and options used for the benchmarks in Fig. 3 are described in Table 3. Their performance is also described in Table 4 (this contains the same information as Fig. 3, but as numbers rather than a bar chart). For a full list of benchmarks carried out, and the options used, see Supplementary Section B. Furthermore, Supplementary Fig. D.1 shows the performance of all trialed combinations of methods and options, with Supplementary Fig. D.2 showing the performance when the simulations are carried out for increasing final model (physical) times.

**Table 3:** Options used for the benchmarked ODE methods displayed in the main text figure. For each model the options used for the 3 most performant native Julia solvers, the Julia lsoda and CVODE implementations, and each other tool (the results using these benchmarks are found in Fig. 3). Each field contains the method used for that model. Further options (including whenever a specific linear solver was selected) are described through superscript tags. ^1^ GMRES linear solver was used. ^2^ Lapack Dense linear solver was used. ^3^ Sparse Jacobian representation was used (a Catalyst option only). ^4^ Automatic differentiation (as a mean of Jacobian calculation) was turned off (a Catalyst option only). ^5^ An incomplete LU precondition was supplied to the GMRES linear solver (a Catalyst option only).

| Model: | Multistate | Multisite2 | Egfr_net | BCR | Fceri_gamma2 |
| --- | --- | --- | --- | --- | --- |
| Julia solver 1 | Vern6 | BS5 | VCABM | QNDF<sup>1,3,4,5</sup> | QNDF<sup>1,3,4,5</sup> |
| Julia solver 2 | Vern7 | Vern8 | BS5 | FBDF<sup>1,3,4,5</sup> | FBDF<sup>1,3,4,5</sup> |
| Julia solver 3 | Tsit5 | Tsit5 | Vern6 | KenCarp4<sup>1,3,4,5</sup> | TRBDF2<sup>1,3,4,5</sup> |
| Catalyst lsoda | lsoda | lsoda | lsoda | lsoda | lsoda |
| Catalyst CVODE | CVODE | CVODE<sup>1</sup> | CVODE<sup>1</sup> | CVODE<sup>1,3,5</sup> | CVODE<sup>1,3,5</sup> |
| BioNetGen | CVODE<sup>1</sup> | CVODE<sup>1</sup> | CVODE<sup>1</sup> | CVODE | CVODE<sup>1</sup> |
| COPASI | CVODE | CVODE | CVODE | CVODE | CVODE |
| GillesPy2 | lsoda | lsoda | lsoda | lsoda | lsoda |
| Matlab | CVODE | CVODE | CVODE | CVODE | CVODE |

**Table 4:** ODE Benchmark times. Fig. 3 A-E shows the results as bar charts, here, the same benchmark times (median over several simulations) are given as numbers (in units of milliseconds). Each field corresponds to the same field in Table 3.

| Model: | Multistate | Multisite2 | Egfr_net | BCR | Fceri_gamma2 |
| --- | --- | --- | --- | --- | --- |
| Julia solver 1 | 0.107 | 0.477 | 5.89 | 9,550 | 1,570 |
| Julia solver 2 | 0.114 | 0.517 | 5.91 | 12,500 | 1,670 |
| Julia solver 3 | 0.117 | 0.535 | 6.65 | 35,900 | 7,510 |
| Catalyst lsoda | 0.220 | 1.17 | 29.7 | 172,000 | 644,000 |
| Catalyst CVODE | 0.310 | 1.78 | 7.31 | 6,520 | 423 |
| BioNetGen | 154 | 322 | 5,280 | 170,000 | 108,000 |
| COPASI | 0.849 | 13.0 | 520 | 3,110,000 | 1,990,000 |
| GillesPy2 | 17.2 | 315 | 6,280 | 24,400,000 | 26,800,000 |
| Matlab | 8.34 | 47.8 | 880 | 101,000 | 354,000 |

Stochastic chemical kinetics simulations of Catalyst models used SSAs defined in JumpProcesses.jl [50], a component of DifferentialEquations.jl. In Figure 3, Direct refers to Gillespie’s direct method [9], SortingDirect to the sorting direct method of [51], RSSA and RSSACR to the rejection and composition-rejection SSA methods of [75–77]. Dependency graphs needed for the different methods are automatically generated via Catalyst and ModelingToolkit as input to the JumpProcesses.jl solvers. Due to supercloud not permitting single runs longer than 4 days, for the largest models, the slowest tools and methods were not benchmarked. The BCR model exhibits pulses, to ensure that at least some pulses were included in each SSA simulation, this model was simualted over very long timespans (*>* 10, 000 seconds). For a full list of SSA benchmarks and their options, please see Supplementary Section C.

The benchmarks were carried out on Julia version 1.8.5, using Catalyst version 13.1.0, JumpProcesses version 9.5.1, and OrdinaryDiffEq version 6.49.0. Note that JumpProcesses and OrdinaryDiffEq are both components in the meta DifferentialEquations.jl package. We used Python version 3.9.15, the version 0.7.9 python interface for BioNetGen, the basico version 4.47 python interface for COPASI, GillesPy2 version 1.8.1, and Matlab version 9.8 with SimBiology version 5.10.

## 5 Code availability

Scripts for generating all figures presented here, as well as for carrying out the benchmarks, can be found at https://github.com/TorkelE/Catalyst-Fast-Biochemical-Modeling-with-Julia.

## 6 Acknowledgements

The authors thank the 26 other individuals who contributed commits to Catalyst, the Catalyst tutorials, and the Catalyst documentation, along with the many users who have offered suggestions and opened issues.

TL’s contribution to this project has received funding from the European Union’s Horizon 2020 research and innovation programme under the Marie Sklodowska-Curie grant agreement No.721456.

SAI’s and CR’s work on this project has been made possible in part by the following two grants to the SciML organization. This research was funded in whole, or in part, by the Wellcome Trust [223770/Z/21/Z]. For the purpose of open access, the author has applied a CC BY public copyright licence to any Author Accepted Manuscript version arising from this submission. This publication and software have been made possible in part by CZI grant DAF2021-237457 and grant DOI https://doi.org/10.37921/149019qvrhgz from the Chan Zuckerberg Initiative DAF, an advised fund of Silicon Valley Community Foundation (funder DOI 10.13039/100014989). SAI was also partially supported by National Science Foundation DMS-1902854. VI was partially supported by a 2021 Google Summer of Code Fellowship and the Boston University UROP program.

CR’s contribution to this material is based upon work supported by the National Science Foundation under grant no. OAC-1835443, grant no. SII-2029670, grant no. ECCS-2029670, grant no. OAC-2103804, and grant no. PHY-2021825. We also gratefully acknowledge the U.S. Agency for International Development through Penn State for grant no. S002283-USAID. The information, data, or work presented herein was funded in part by the Advanced Research Projects Agency-Energy (ARPA-E), U.S. Department of Energy, under Award Number DE-AR0001211 and DE-AR0001222. This material is based upon work supported by the Defense Advanced Research Projects Agency (DARPA) under Agreement No HR00112290091. We also gratefully acknowledge the U.S. Agency for International Development through Penn State for grant no. S002283-USAID. The views and opinions of authors expressed herein do not necessarily state or reflect those of the United States Government or any agency thereof. This material was supported by The Research Council of Norway and Equinor ASA through Research Council project “308817 - Digital wells for optimal production and drainage”. Research was sponsored by the United States Air Force Research Laboratory and the United States Air Force Artificial Intelligence Accelerator and was accomplished under Cooperative Agreement Number FA8750-19-2-1000. The views and conclusions contained in this document are those of the authors and should not be interpreted as representing the official policies, either expressed or implied, of the United States Air Force or the U.S. Government. The U.S. Government is authorized to reproduce and distribute reprints for Government purposes notwithstanding any copyright notation herein.

The authors acknowledge the MIT SuperCloud and Lincoln Laboratory Supercomputing Center for providing (HPC, database, consultation) resources that have contributed to the research results reported within this paper/report.

## 7 Author contributions

CR, SAI, and TEL conceptualized and developed Catalyst, with NY extending Catalyst’s network analysis functionality. NK aided in the early DSL conceptualization, and developed the Catalyst-Latexify bindings for Latex printing. CR, SG, and YM conceptualized and developed ModelingToolkit and Symbolics, with SAI contributing Catalyst and jump process-related functionality to ModelingToolkit. CR, SAI, and VI conceptualized and developed the SSAs in JumpProcesses (and DifferentialEquations). CR and YM, conceptualized and developed much of the ODE and SDE solver functionality in DifferentialEquations. CR, SAI and TEL conceptualized the simulation benchmarks. TEL performed the simulation benchmarks. CR, SAI, and TEL wrote the manuscript.

## A Exact ODE benchmark times

This section contains Supplementary Table 4, describing the exact benchmark times plotted in Figure 3 A-E.

## B List of ODE benchmarks

Here follows a list of all ODE simulation methods, and combinations of options, used for each tool. The best-performing combinations are shown in Fig. 3 (with details of the used options listed in Table 3). The results of all combinations listed here can be found in Supplementary Figs. D.1 and D.2. For some settings, a parenthesis further clarifies what the option implies. For all ODE simulations, the absolute tolerance was set to `1e-9` and the relative tolerance to `1e-6`.

### B.1 BioNetGen solvers

The `ode` method is the CVODE solver, by default using dense LU and a finite difference Jacobian approximation. The `n steps` parameter sets the number of saved steps. We sat this to 1, to ensure no intermediary time points were saved. Setting `sparse` = `true` enables the (un-preconditioned) GMRES linear solver (for which we used the default GMRES tolerance). BioNetGen does not seem to offer an option to specify a preconditioner for GMRES.

- `Method` = `ode`, `n steps` = `1`, `sparse` = `false`.
- `Method` = `ode`, `n steps` = 1, `sparse` = `true`.

### B.2 Catalyst solvers

The `CVODE_BDF` sovler is the CVODE solver. The `KrylovJL_GMRES` linear solver is the GMRES linear solver (for which we used the default GMRES tolerance). The `jac` = `true` options set that a symbolic Jacobian is built, else, the Jacobian is computed through automatic differentiation (or if `autodiff` = `false` is used, through finite differences). If the `sparse` = `true` is used, a sparse Jacobian representation is used.

- Solver = `lsoda`.
- Solver = `CVODE_BDF`.
- Solver = `CVODE_BDF`, `linear solver` = `LapackDense`.
- Solver = `CVODE_BDF`, `linear solver` = `GMRES`.
- Solver = `CVODE_BDF`, `linear solver` = `GMRES`, `sparse` = `true`, iLU preconditioner.
- Solver = `CVODE_BDF`, `linear solver` = `KLU`, `sparse` = `true`, `jac` = `true`.
- Solver = `TRBDF2`.
- Solver = `TRBDF2`, `linsolve` = `KrylovJL_GMRES`.
- Solver = `TRBDF2`, `linsolve` = `KrylovJL_GMRES`, `sparse` = `true`, iLU preconditioner.
- Solver = `TRBDF2`, `linsolve` = `KLUFactorization`, `sparse` = `true`, `jac` = `true`.
- Solver = `KenCarp4`.
- Solver = `KenCarp4`, `linsolve` = `KrylovJL_GMRES`.
- Solver = `KenCarp4`, `linsolve` = `KrylovJL_GMRES`, `sparse` = `true`, iLU preconditioner.
- Solver = `KenCarp4`, `linsolve` = `KLUFactorization`, `sparse` = `true`, `jac` = `true`.
- Solver = `QNDF`.
- Solver = `QNDF`, `linsolve` = `KrylovJL_GMRES`.
- Solver = `QNDF`, `linsolve` = `KrylovJL_GMRES`, `sparse` = `true`, iLU preconditioner.
- Solver = `QNDF`, `linsolve` = `KLUFactorization`, `sparse` = `true`, `jac` = `true`.
- Solver = `FBDF`.
- Solver = `FBDF`, `linsolve` = `KrylovJL_GMRES`.
- Solver = `FBDF`, `linsolve` = `KrylovJL_GMRES`, `sparse` = `true`, iLU preconditioner.
- Solver = `FBDF`, `linsolve` = `KLUFactorization`, `sparse` = `true`, `jac` = `true`.
- Solver = `Rodas4`.
- Solver = `Rodas4`, `linsolve` = `KrylovJL_GMRES`.
- Solver = `Rodas4`, `linsolve` = `KrylovJL_GMRES`, `sparse` = `true`, iLU preconditioner.
- Solver = `Rodas4`, `linsolve` = `KLUFactorization`, `sparse` = `true`, `jac` = `true`.
- Solver = `Rodas5P`.
- Solver = `Rodas5P`, `linsolve` = `KrylovJL_GMRES`.
- Solver = `Rodas5P`, `linsolve` = `KrylovJL_GMRES`, `sparse` = `true`, iLU preconditioner.
- Solver = `Rodas5P`, `linsolve` = `KLUFactorization`, `sparse` = `true`, `jac` = `true`.
- Solver = `Rosenbrock23`.
- Solver = `Rosenbrock23`, `linsolve` = `KrylovJL_GMRES`.
- Solver = `Rosenbrock23`, `linsolve` = `KrylovJL_GMRES`, `sparse` = `true`, iLU preconditioner.
- Solver = `Rosenbrock23`, `linsolve` = `KLUFactorization`, `sparse` = `true`, `jac` = `true`.
- Solver = `Tsit5`.
- Solver = `BS5`.
- Solver = `VCABM`.
- Solver = `Vern6`.
- Solver = `Vern7`.
- Solver = `Vern8`.
- Solver = `Vern9`.
- Solver = `ROCK2`.
- Solver = `ROCK4`.

The iLU preconditioner option simulations were benchmarked only for the two largest models (BCR and Fceri gamma2). The explicit solvers (`Tsit5`, `BS5`, `VCABM`, `Vern6`, `Vern7`, `Vern8`, `Vern9`, `ROCK2`, and `ROCK4`) were not benchmarked for the BCR model.

For all benchmarks, we used the `saveat` = [Simulation length] option to ensure no intermediary time points were saved. For the `TRBDF2`, `KenCarp4`, `QNDF`, `FBDF`, `Rodas4`, `Rodas5P`, `Rosenbrock23` methods, when benchmarked on the (large) BCR and Fceri gamma2 models, the `autodiff` = `false` option was used to disable automatic differentiation. The iLU preconditioner requires setting a single parameter *τ*. For CVODE we used *τ* = 5 (BCR model) and *τ* = 1*e*2 (Fceri gamma2 model). For the other methods, we used *τ* = 1*e*12 (BCR model) and *τ* = 1*e*2 (Fceri gamma2 model). While these *τ* values were roughly optimized for their specific problems, we note that for both the BCR and Fceri gamma2 models, the fastest Catalyst solvers included solvers not depending on preconditioners with customized *τ* values.

### B.3 COPASI solvers

The `stepsize` option sets the time between saving the solution. We used the length of the simulation to ensure no intermediary time points were saved.

- `method` = `deterministic` (CVODE), `stepsize` = [Simulation length].

### B.4 GillesPy2 solvers

The `nsteps` option sets the maximum number of time steps. The `increment` option sets the time between saving the solution. We used the length of the simulation to ensure no intermediary time points were saved. The `solver` = `ODESolver` options designate that we run ODE simulations (as opposed to e.g. Gillespie simulations), while the `integrator` sets the numerical solver method used (e.g. lsoda).

- `solver` = `ODESolver`, `integrator` = `lsoda`, `nsteps` = `100000`, `increment` = [Simulation length].
- `solver` = `ODESolver`, `integrator` = `csolver`, `nsteps` = 100000, `increment` = [Simulation length].
- `solver` = `ODESolver`, `integrator` = `zvode`, `nsteps` = `100000`, `increment` = [Simulation length].
- `solver` = `ODESolver`, `integrator` = `vode`, `nsteps` = `100000`, `increment` = [Simulation length].

Due to their poor performance compared to `lsoda`, the `zvode` and `vode` methods benchmarks were investigated beyond initial tests. Furthermore, due to its poor performance, the `csolver` solver was not benchmarked for the two largest (BCR and Fceri gamma2) models.

### B.5 Matlab solvers

The `sundials` solver type is the CVODE solver, using its default options. For Matlab there appeared to be no other documented control parameters (including setting the density at which time points were saved) that could beneficially affect the performance. We tried the `RuntimeOptions.StatesToLog =` *{}* option, however, this offered no improvement, while reducing performance for the smaller models.

- `SolverType` = `sundials`.

## C List of SSA benchmarks

Here follows a list of all SSA simulation methods used for each tool. Their performance is shown in Fig. 3 (due to the small number of SSA methods, all fit in Fig. 3 and no supplementary figure was needed).

### C.1 BioNetGen solvers

The `ssa` method is the Sorting Direct method. The `n steps` parameter sets the number of saved steps. We sat this to 1, to ensure no intermediary time points were saved.

- `method` = `ssa`, `n steps` = `1`.

Due to the benchmarks surpassing the 4-day limit of jobs at the used HPC supercomputer, the `ssa` method was not bench-marked on the BCR model.

### C.2 Catalyst solvers

Here, the `save positions` = `(false,false)` options prevent the saving of the solution before/after a jump is made (which will severely reduce performance when a large number of jumps are performed). Using this option, we ensured that no intermediary time points were saved.

- Solver = `Direct`, `save positions` = `(false,false)`
- Solver = `SortingDirect`, `save positions` = `(false,false)`
- Solver = `RSSA`, `save positions` = `(false,false)`
- Solver = `RSSACR`, `save positions` = `(false,false)`

### C.3 Copasi solvers

- `method` =`directMethod`, `stepsize` = [Simulation length].

Due to the benchmarks surpassing the 4-day limit of jobs at the used HPC supercomputer, COPASI was not benchmarked on the BCR model.

### C.4 GillesPy2 solvers

The `nsteps` option sets the maximum number of time steps. The `increment` option sets the time between saving the solution. We used the length of the simulation to ensure no intermediary time points were saved.

- `solver` = `ssa`, `increment` = [Simulation length].
- `solver` = `numpyssa`, `increment` = [Simulation length].

Due to its poor performance compared to `ssa`, the `numpyssa` method benchmarks were never concluded beyond initial tests. Due to the benchmarks surpassing the 4-day limit of jobs at the used HPC supercomputer, the `ssa` method was not benchmarked on the BCR and Fceri gamma2 models.

### C.5 Matlab solvers

The `ssa` solver type is Gillespie’s Direct method. For Matlab there appeared to be no other documented control parameters (including setting the density at which time points were saved) that could affect the performance.

- `SolverType` = `ssa`.

Due to slow performance and/or crashing for larger models, the `ssa` method was not benchmarked on the BCR and Fceri gamma2 models.

## D Additional benchmarks of ODE solvers

In this article we compare ODE simulation benchmarks for Catalyst and various other CRN modelling tools (BioNetGen, COPASI, GillesPy2, and Matlab’s SimBiology toolbox) (Fig. 3). While we benchmarked a large number of combinations of methods and tools (Supplementary Section B), the main text figure only displays the results using the best combinations. Here, in Supplementary Figs. D.1 and D.2, we show the benchmarks for the full set. Supplementary Fig. D.1 shows the run time to reach each models’ steady states (or, for the BCR model, complete 3 pulses), while Supplementary Fig. D.2 shows the simulation times as a function of the real (physical) end time of the simulation. The exact options used for each simulation are described in Supplementary Section B.

**Figure D.1:**
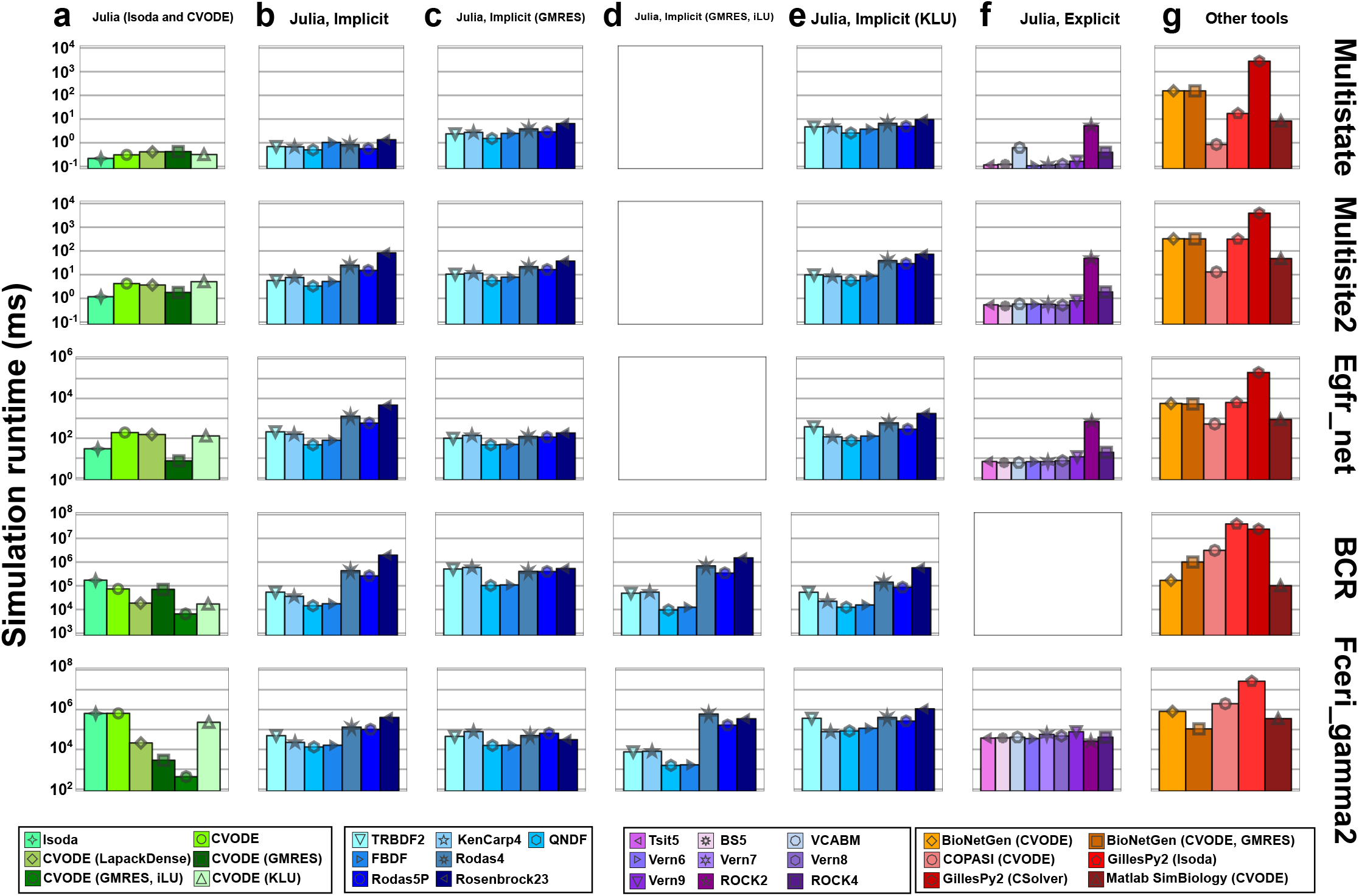
The result of the benchmarks across the full set of ODE solvers and options. These benchmarks are a more extensive set than what is provided in Fig. 3 (which includes only the best alternative for each combination of tool and model). (a) Benchmarks for Catalyst, using lsoda, as well as CVODE with a range of options (no linear solver specified, or for CVODE, the LapackDense, GMRES, or KLU linear solvers). The GMRES option was trialed with and without the iLU preconditioner. A sparse Jacobian representation was used when the KLU linear solver or iLU preconditioner was used. Furthermore, for the KLU linear solver a symbolic, sparse, Jacobian was used. (b) Catalyst benchmarks using an extended set of (implicit) ODE solvers. (c) Benchmark for the same solvers as in B, but with the GMRES linear solver. (d) Same as in C, but using an iLU preconditioner and sparse finite-difference Jacobian representation. (e) Benchmark for the same solvers as in B, but with the KLU linear solver and sparse, symbolic, Jacobian representation. (f) Catalyst benchmarks using an extended set of (explicit) ODE solvers. (g) Benchmarks using non-Catalyst tools. For more details on the options used for each benchmark, please see Supplementary Section B.

**Figure D.2:**
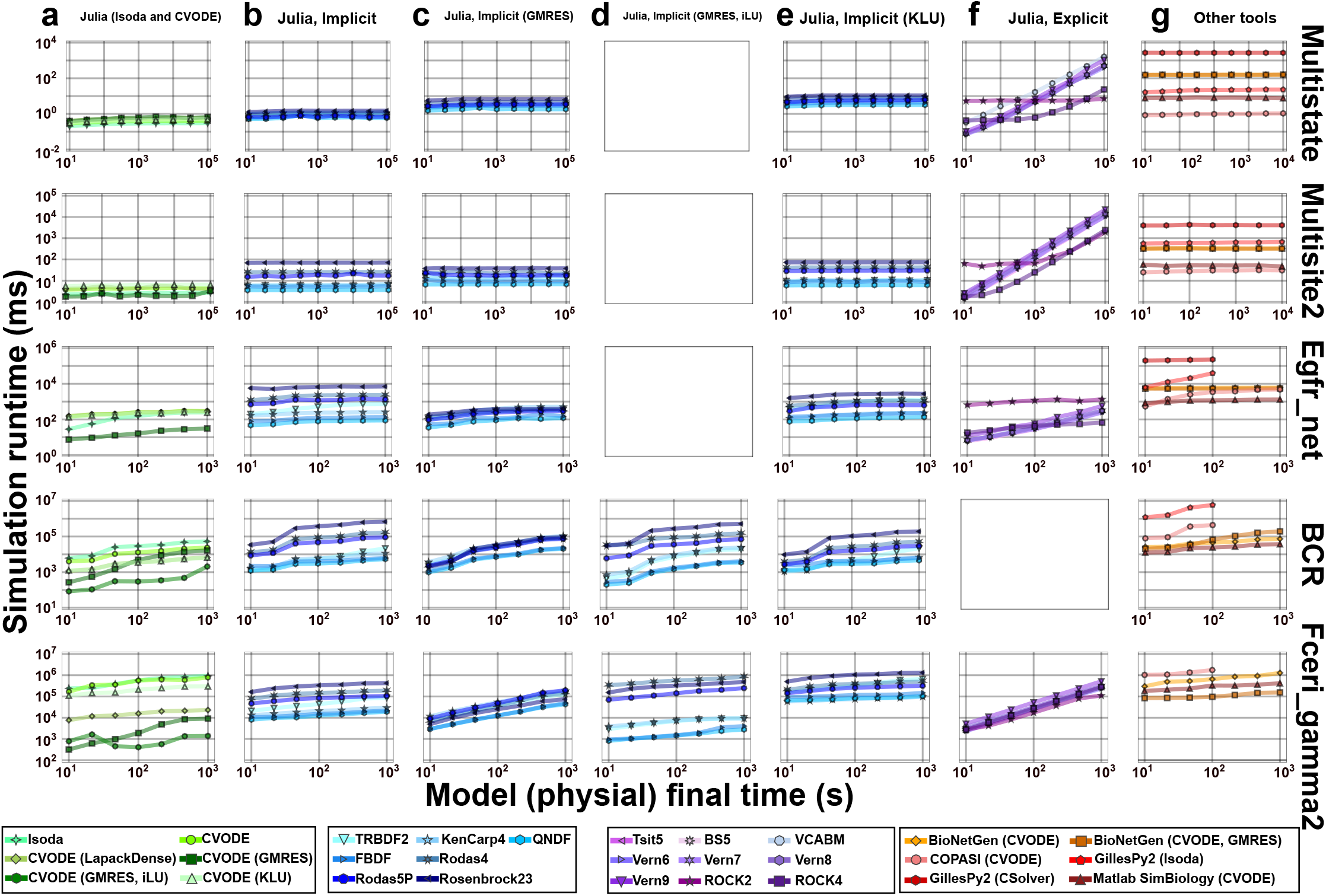
The result of the benchmarks across the full set of ODE solvers and options. These benchmarks are a more extensive set than what is provided in Fig. 3 (which includes only the best alternative for each combination of tool and model). Here, the simulation run time was as a function of the real (physical) end time of the simulation. (a) Benchmarks for Catalyst, using lsoda, as well as CVODE with a range of options (no linear solver specified, or for CVODE, the LapackDense, GMRES, or KLU linear solvers). The GMRES option was trialed using with and without the iLU precondition. A sparse Jacobian representation was used when the KLU linear solver or iLU preconditioner was used. Furthermore, for KLU linear solver, a symbolic, sparse, Jacobian was used. (b) Catalyst benchmarks using an extended set of (implicit) ODE solvers. (c) Benchmark for the same solvers as in B, but with the GMRES linear solver. (d) Same as in C, but using an iLU preconditioner and a sparse finite-difference Jacobian representation. (e) Benchmark for the same solvers as in B, but with the KLU linear solver and sparse, symbolic, Jacobian representation. (f) Catalyst benchmarks using an extended set of (explicit) ODE solvers. (g) Benchmarks using non-Catalyst tools. For more details on the options used for each benchmark, please see Supplementary Section B.

## E Work-precision diagrams for Julia solvers

For the best-performing Julia solvers, we compare the run time of the numerical simulator to the error it generates. For each combination of solver and options, we make repeated simulations at various tolerances (absolute and real tolerance both equal to 10^*−*5^, 10^*−*6^, 10^*−*7^, or 10^*−*8^). For each simulation, we calculate the error by comparison to a single simulation using the CVODE solver and no options, with absolute and real tolerances for the latter equal to 10^*−*12^. For each tolerance, we can thus compute the mean error and simulation time (across all simulations at that tolerance). Plotting these, we generate a so-called work-precision diagram. Such diagrams for all models are shown in Supplementary Fig. E.1.

Unlike all other benchmarks, the work-precision diagrams were not computed on supercloud, but rather on the SciML organization’s benchmarking server, which has the following specifications:

**OS** Linux (x86_64-linux-gnu)

**CPU** 128 × AMD EPYC 7502 32-Core Processor

**WORD_SIZE** 64

**LIBM** libopenlibm

**LLVM** libLLVM-13.0.1 (ORCJIT, znver2)

**Threads** 128 on 128 virtual cores

The code that was used to generate Supplementary Fig. E.1 can be found in the associated Github benchmarking repository for the article, see Section 5. For a more extensive set of work-precision diagrams of available Julia (and thus Catalyst) solvers, please refer to the SciMLBenchmarks.jl package.

**Figure E.1:**
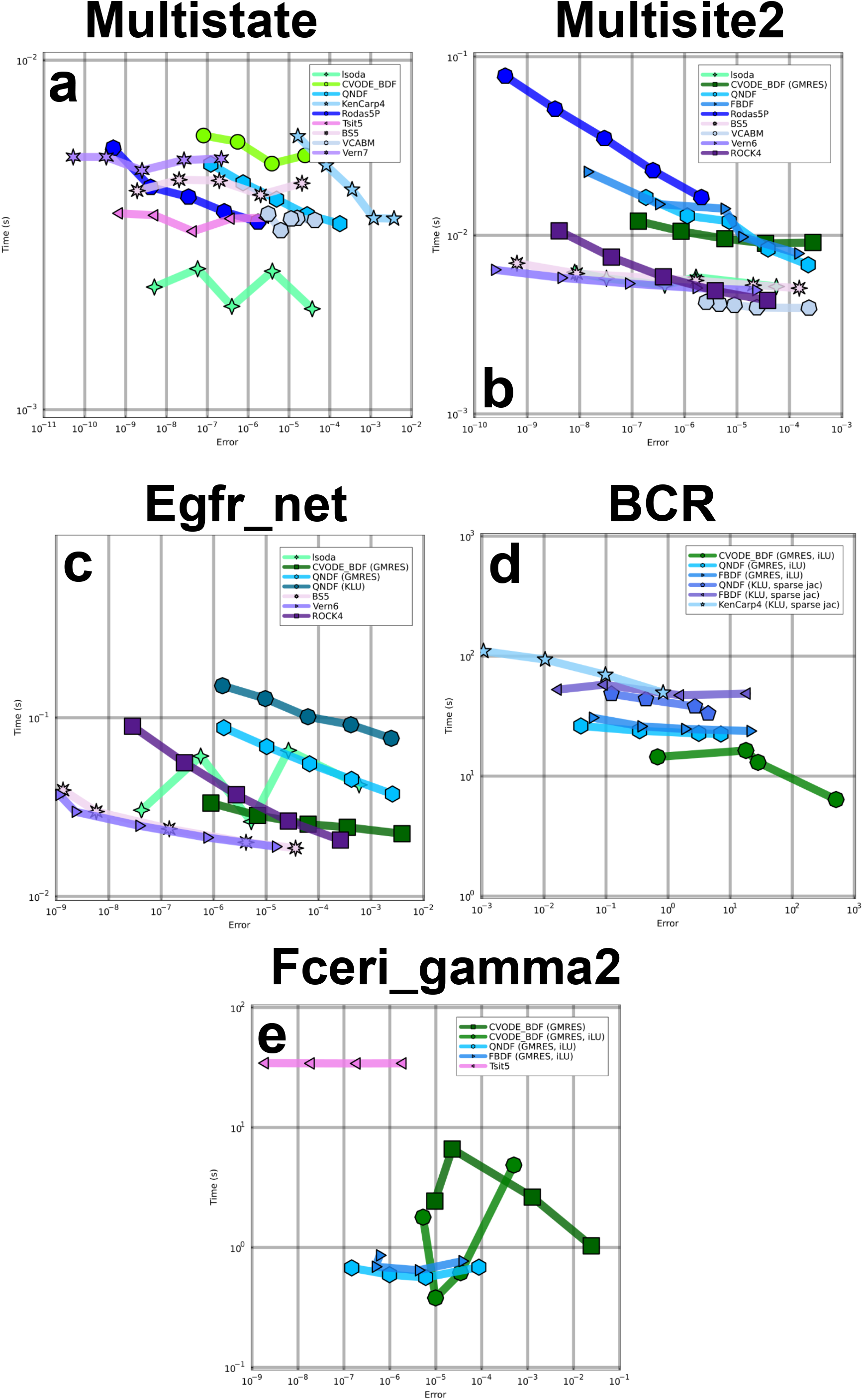
Work-precision diagrams for selected Julia ODE solvers (and options). Native Julia solvers typically have smaller errors (compared to lsoda and CVODE) when tolerances are kept identical. The simulation options used for this figure are drawn from the list in Supplementary Section B.

## F Simulation trajectories for various modelling tools

In this article, we compare ODE and SSA simulation benchmarks for Catalyst and various other CRN modelling tools (BioNet-Gen, COPASI, GillesPy2, and Matlab’s SimBiology toolbox). To ensure that each tool successfully simulates each model, we here plot the output trajectories for each combination of tool and model. These plots are displayed in Supplementary Figs. F.1-F.10.

**Figure F.1:**
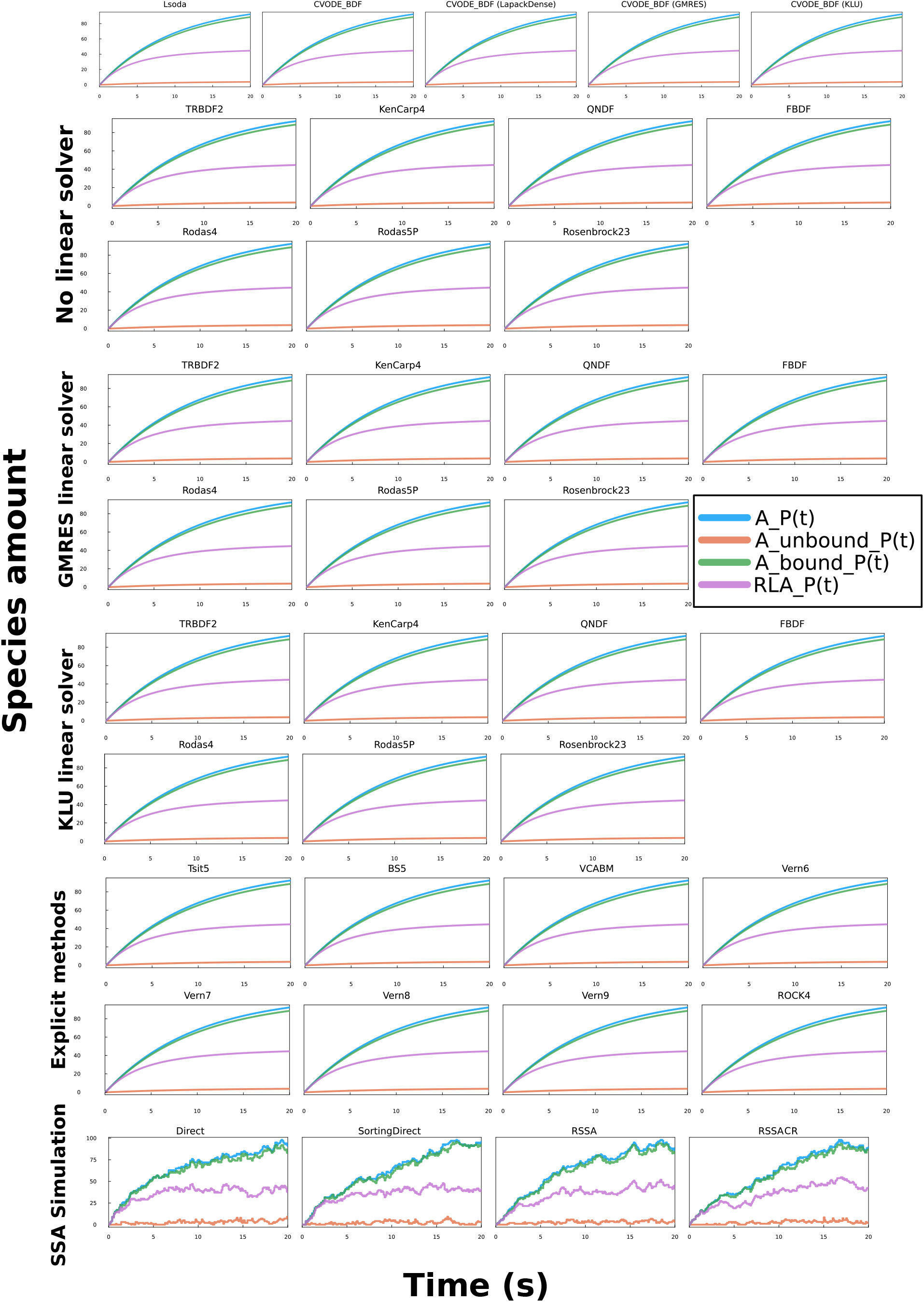
Catalyst benchmark simulation time trajectories for the multistate model. The multistate model is simulated until it reaches its (approximate) steady state at *t* = 20 seconds (same time point which was used for the benchmarks in Fig. 3). It was simulated using both ODE and SSA methods. The simulation trajectories correspond to those of the other tools, suggesting that the models are correctly interpreted.

**Figure F.2:**
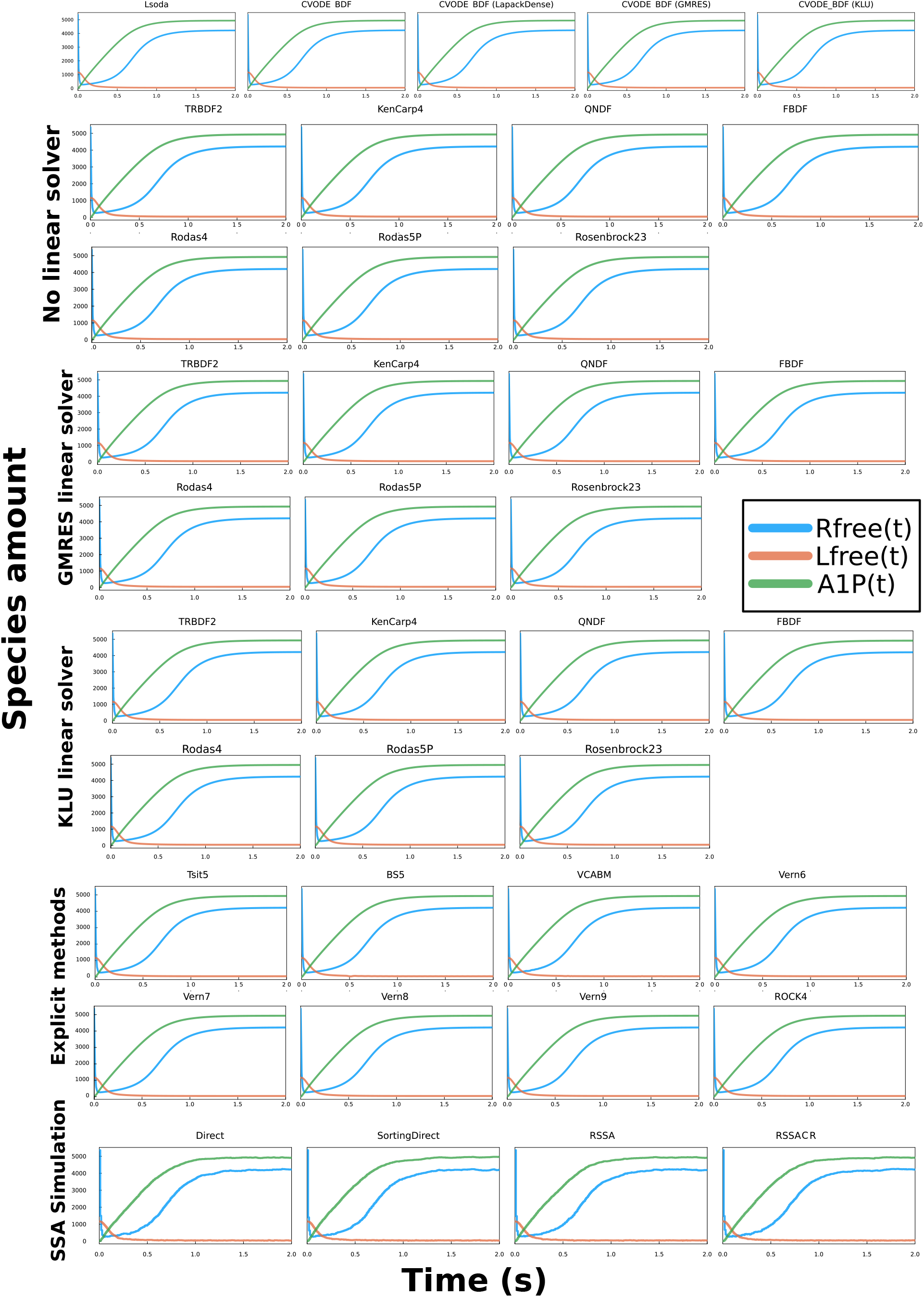
Catalyst benchmark simulation time trajectories for the multisite2 model. The multisite2 model is simulated until it reaches its (approximate) steady state at *t* = 2 seconds (same time point which was used for the benchmarks in Fig. 3). It was simulated using both ODE and SSA methods. The simulation trajectories correspond to those of the other tools, suggesting that the models are correctly interpreted.

**Figure F.3:**
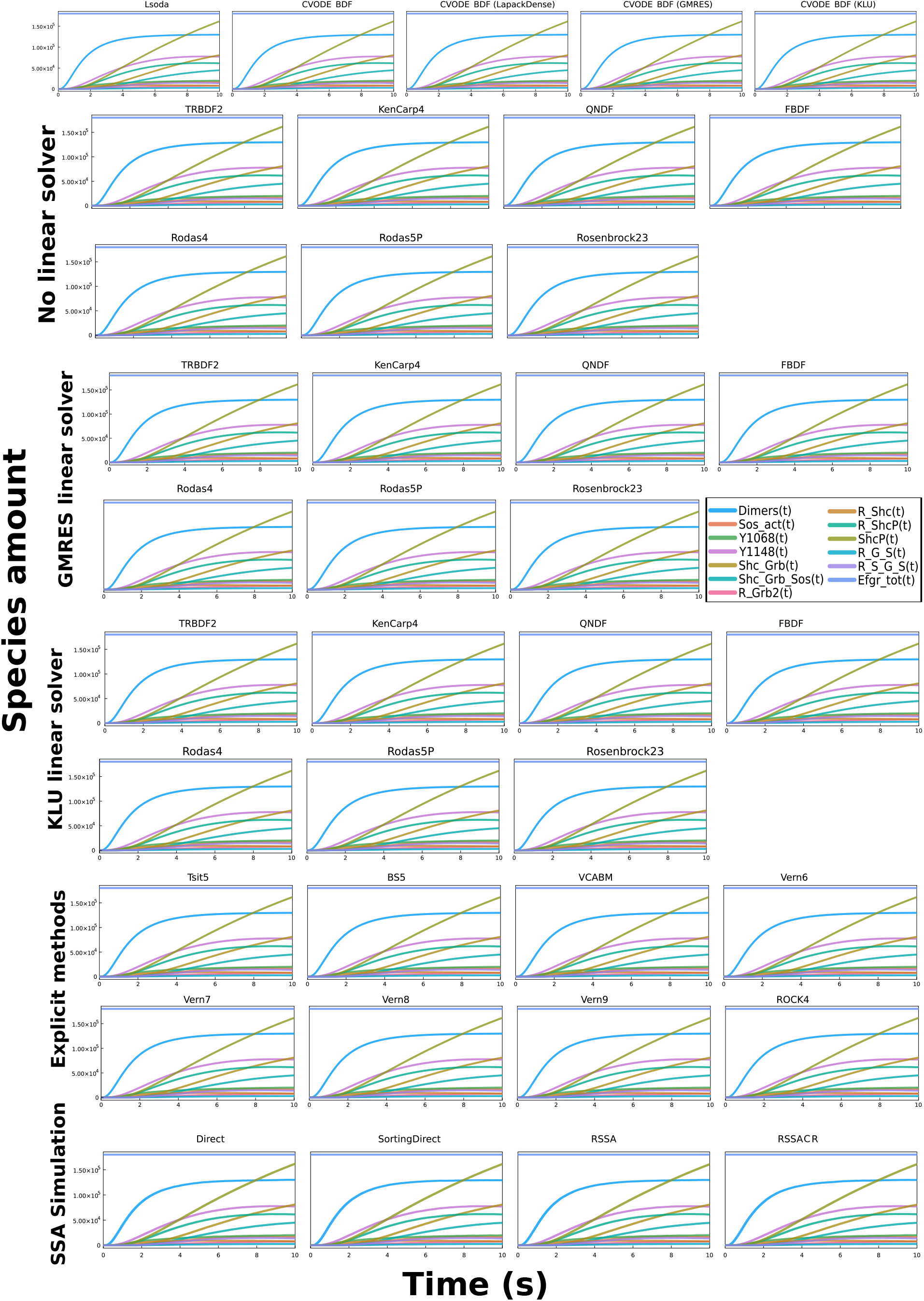
Catalyst benchmark simulation time trajectories for the egfr_net model. The egfr_net model is simulated until it reaches its (approximate) steady state at *t* = 10 seconds (same time point which was used for the benchmarks in Fig. 3). It was simulated using both ODE and SSA methods. The simulation trajectories correspond to those of the other tools, suggesting that the models are correctly interpreted.

**Figure F.4:**
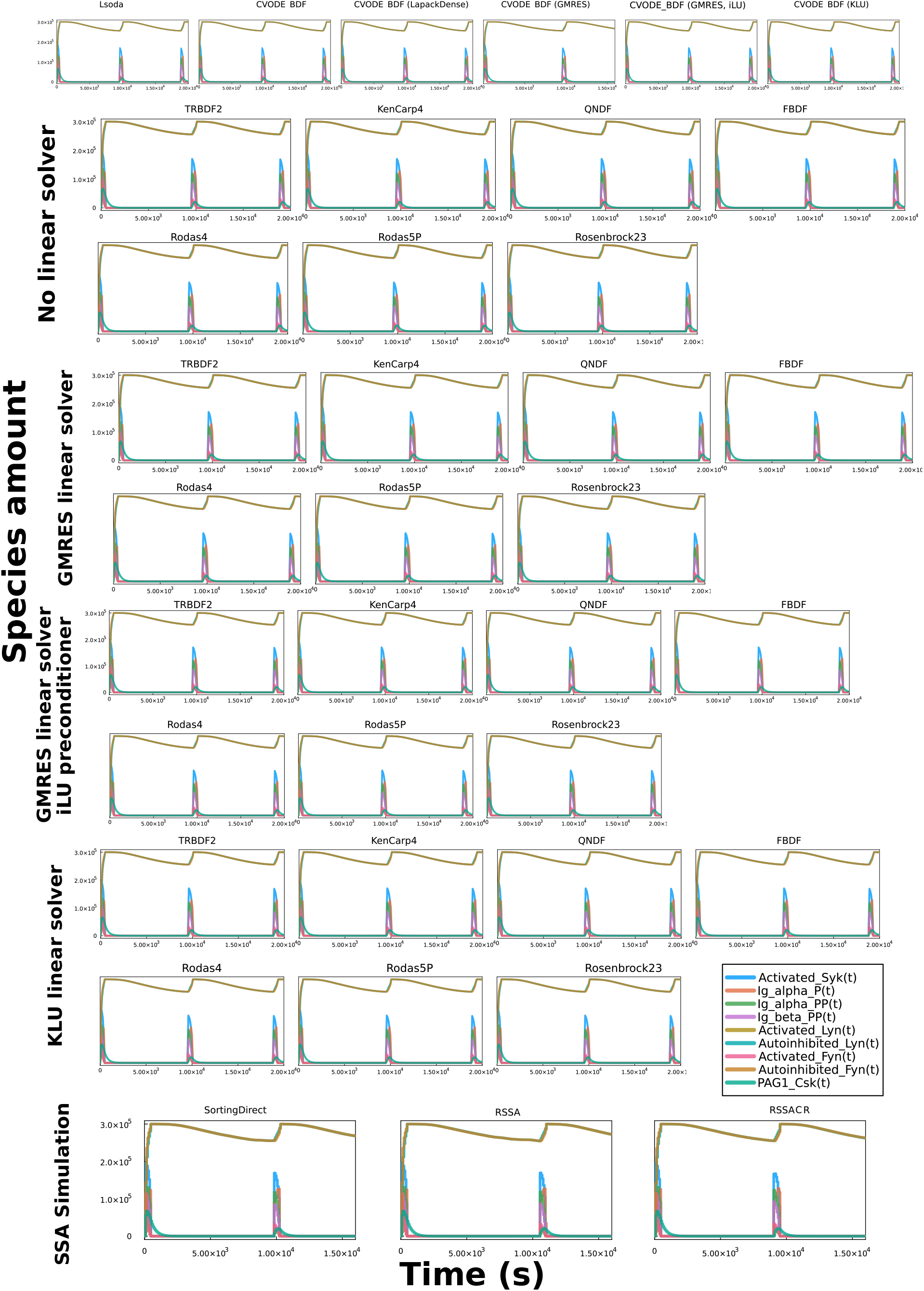
Catalyst benchmark simulation time trajectories for the BCR model. The BCR model is simulated until it reaches its (approximate) steady state at *t* = 10, 000 seconds (same time point which was used for the benchmarks in Fig. 3). It was simulated using both ODE and SSA methods. The time of the SSA simulation’s pulse initiation is variable, and hence the system was resimulated to ensure that a pulse was initiated in the simulation. The simulation trajectories correspond to those of the other tools, suggesting that the models are correctly interpreted.

**Figure F.5:**
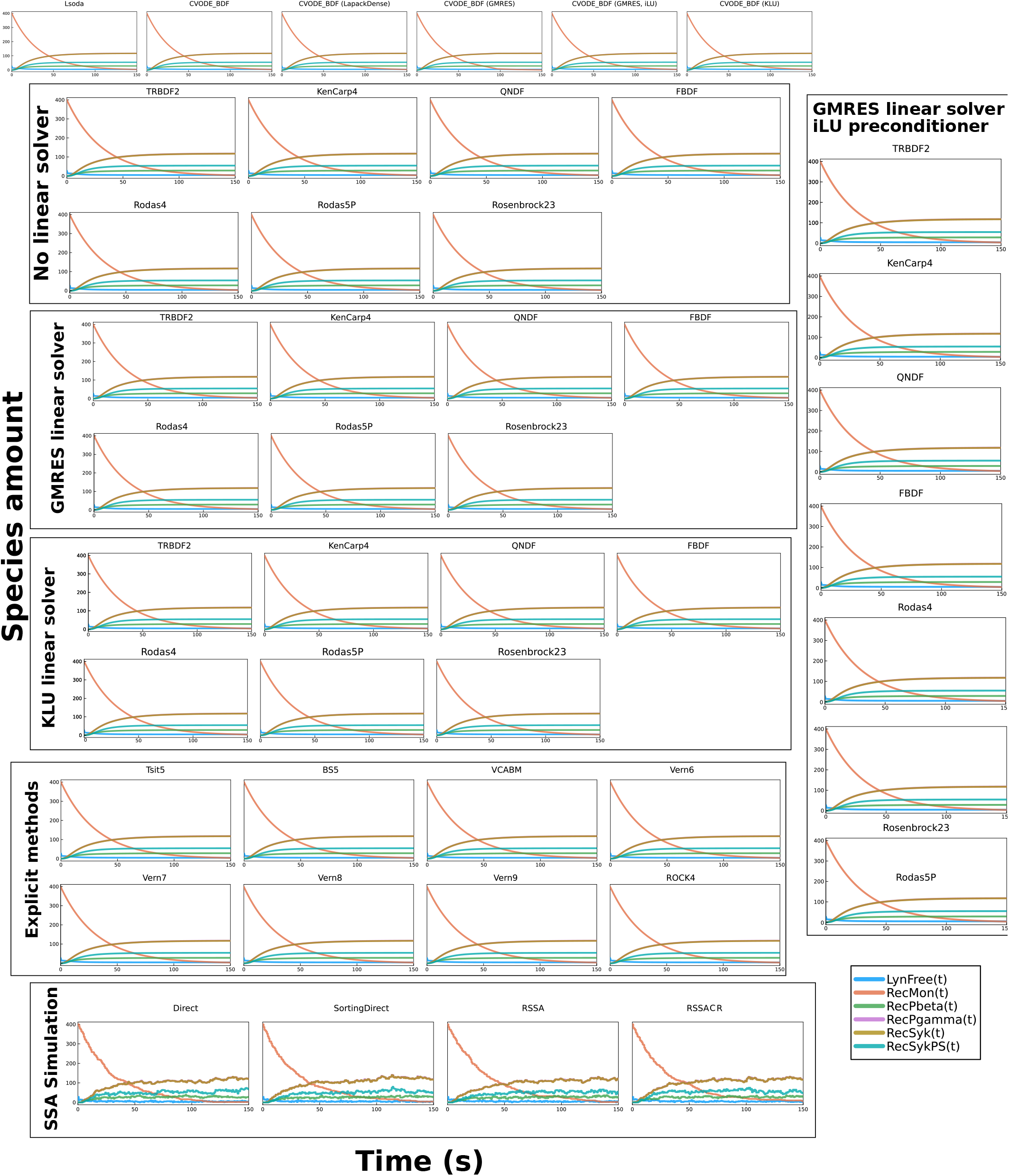
Catalyst benchmark simulation time trajectories for the fceri gamma2 model. The fceri gamma2 model is simulated until it reaches its (approximate) steady state at *t* = 150 seconds (same time point which was used for the benchmarks in Fig. 3). It was simulated using both ODE and SSA methods. The simulation trajectories correspond to those of the other tools, suggesting that the models are correctly interpreted.

**Figure F.6:**
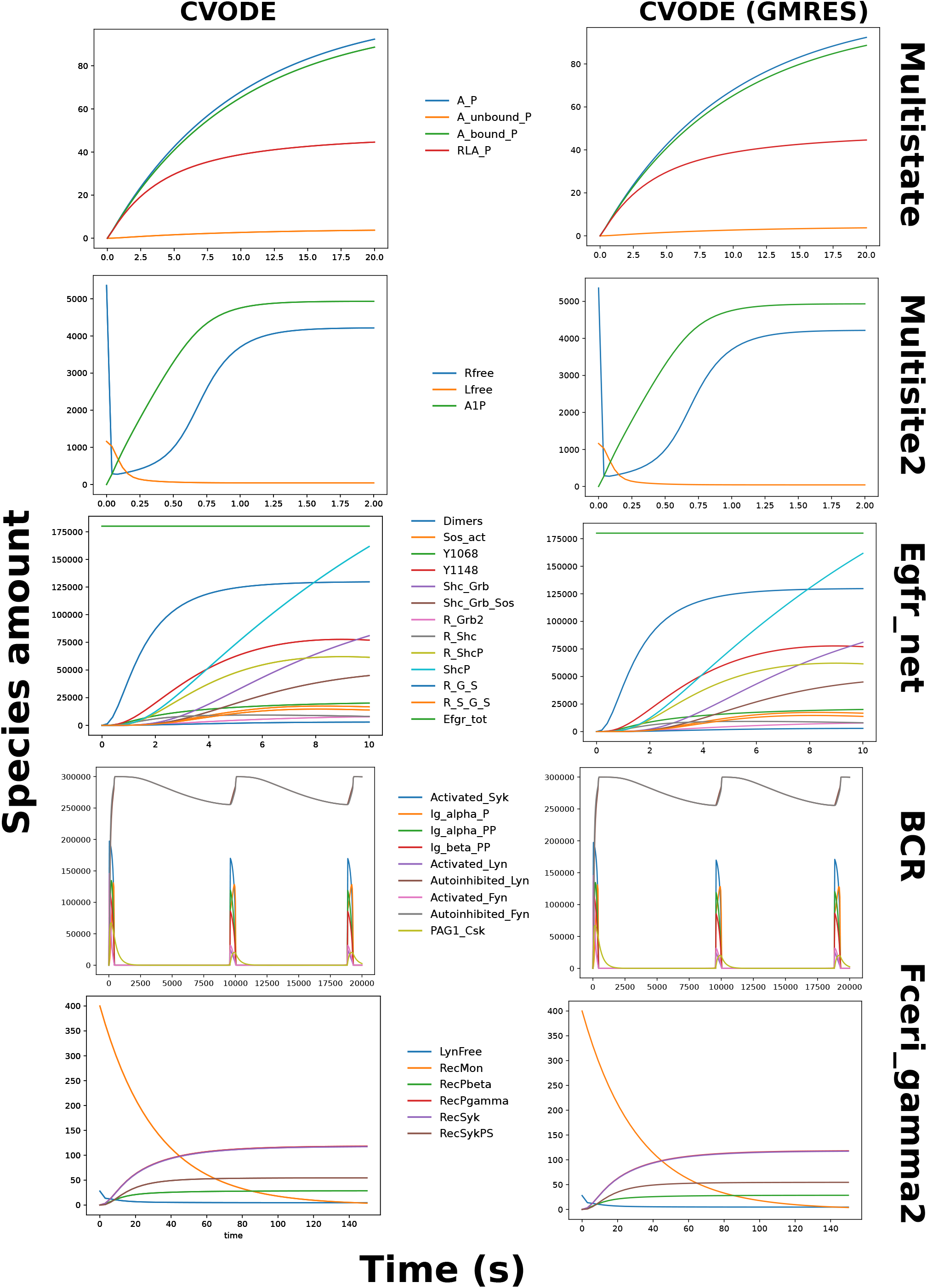
BioNetGen benchmark simulation time trajectories. All models are simulated until they reach their (approximate) steady state (same time point which was used for the benchmarks in Fig. 3). They were simulated for the CVODE method both with and without the GMRES linear solver. The simulation trajectories correspond to those of the other tools, suggesting that the models are correctly interpreted.

**Figure F.7:**
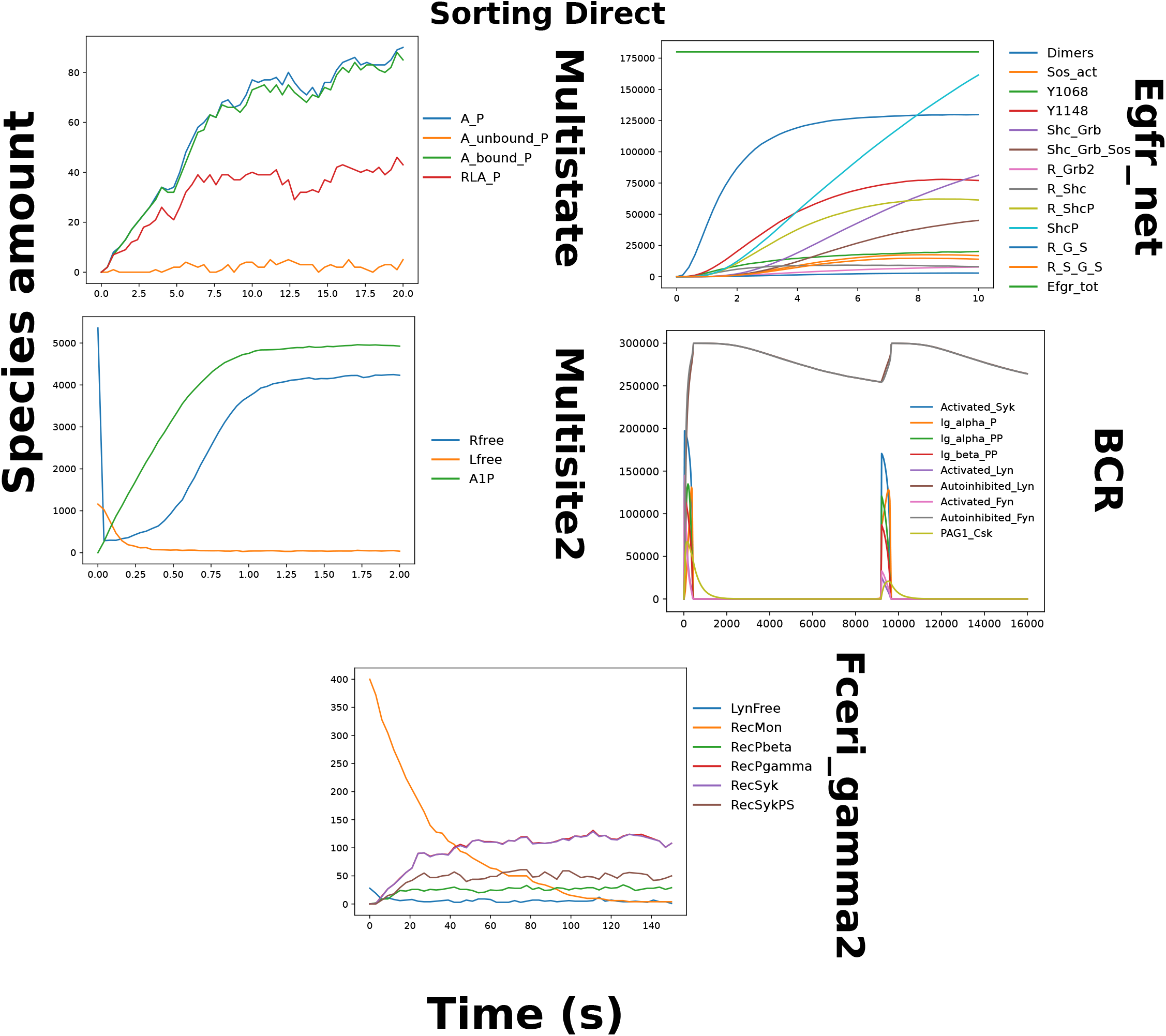
BioNetGen benchmark simulation time trajectories. All models are simulated until they reach their (approximate) steady state (same time point which was used for the benchmarks in Fig. 3). They were simulated using the Sorting Direct method. Due to the long simulation time, we did not produce trajectories for the BCR model. The simulation trajectories correspond to those of the other tools, suggesting that the models are correctly interpreted. Note that the timescale is different for the ODE and SSA simulations.

**Figure F.8:**
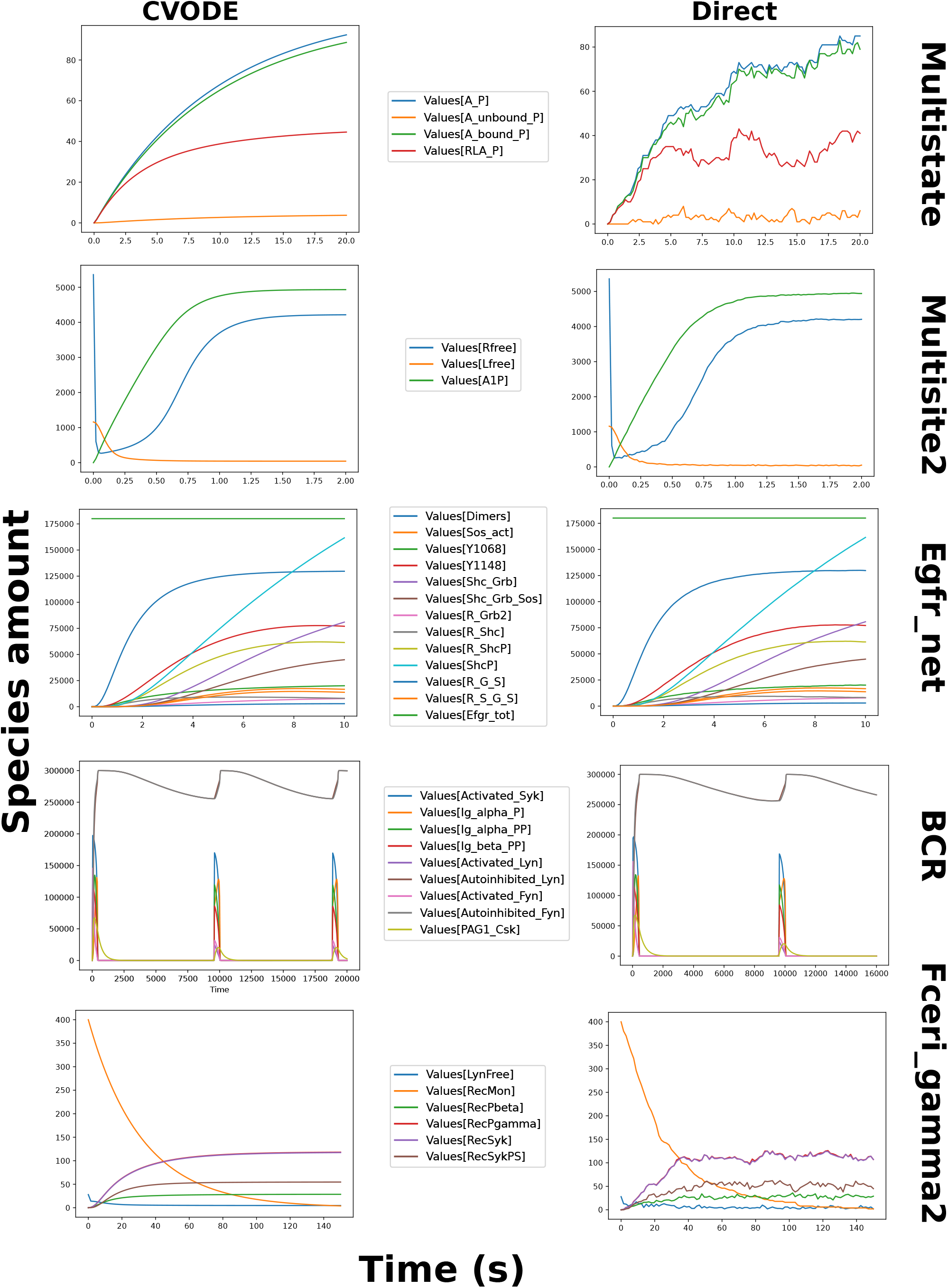
COPASI benchmark simulation time trajectories. All models are simulated until they reach their (approximate) steady state (same time point which was used for the benchmarks in Fig. 3). The simulation trajectories correspond to those of the other tools, suggesting that the models are correctly interpreted.

**Figure F.9:**
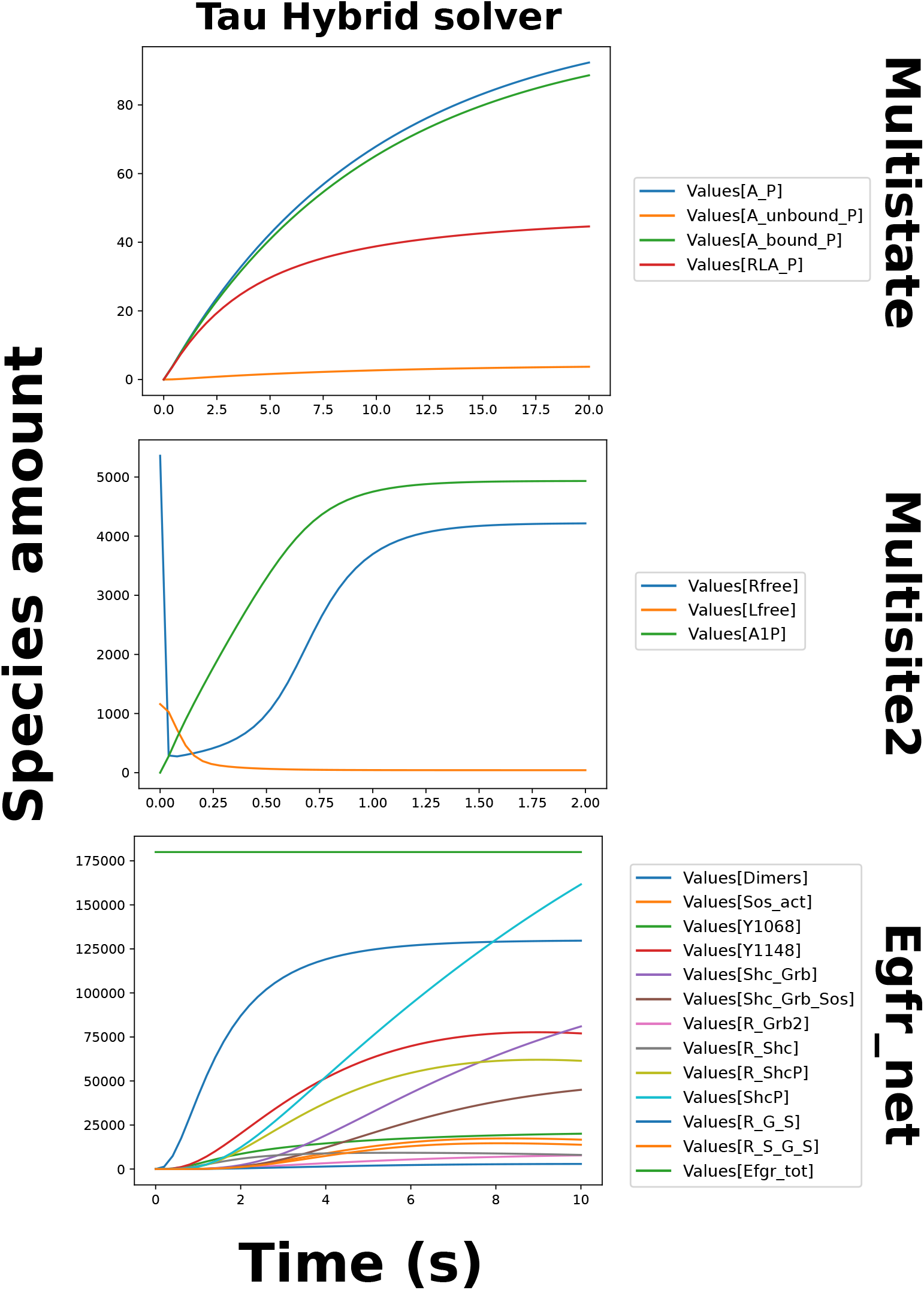
GillesPy2 benchmark simulation time trajectories. All models are simulated until they reach their (approximate) steady state (same time point which was used for the benchmarks in Fig. 3). At the time of investigation, GillesPy2 only permitted the plotting of observables when the Tau hybrid solver was used for simulation. Hence, trajectories could only be checked for this algorithm. Due to the long simulation time required for this method, we were unable to produce trajectories for the two largest models. The simulation trajectories correspond to those of the other tools, suggesting that the models are correctly interpreted.

**Figure F.10:**
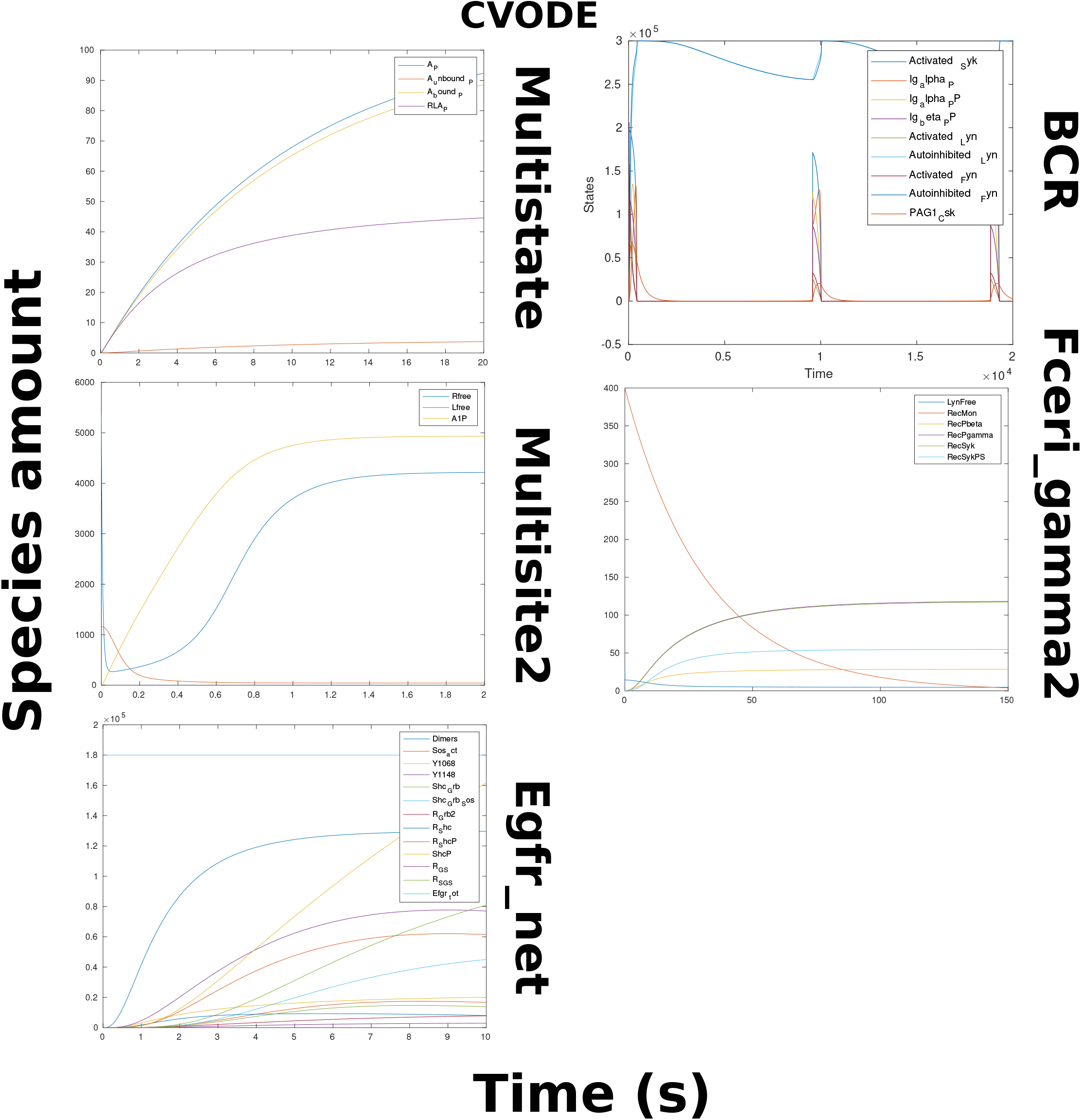
Matlab benchmark simulation time trajectories. All models are simulated until they reach their (approximate) steady state (same time point which was used for the benchmarks in Fig. 3). At the time of investigation, Matlab did not support the plotting of SBML observables from simulations using the Gillespie interpretation, hence we were unable to produce such plots. However, the ODE simulation trajectories correspond to those of the other tools, suggesting that the models are correctly interpreted. Finally, in practice, Matlab was only able to successfully complete Gillespie simulations for the smallest models.

## References

[1] Martin Feinberg. Chemical reaction network structure and the stability of complex isothermal reactors—i. the deficiency zero and deficiency one theorems. Chemical Engineering Science, 42:2229–2268, 1987.

[2] D. Chandran, W.B. Copeland, S.C. Sleight, and H.M. Sauro. Mathematical modeling and synthetic biology. Drug discovery today. Disease models, 5(4):299–309, 2008.

[3] Haluk Resat, Linda Petzold, and Michel F. Pettigrew. Kinetic Modeling of Biological Systems. In Reneé Ireton, Kristina Montgomery, Roger Bumgarner, Ram Samudrala, and Jason McDermott, editors, Computational Systems Biology, Methods in Molecular Biology, pages 311–335. Humana Press, Totowa, NJ, 2009.

[4] Eun-Young Mun and Feng Geng. An epidemic model for non-first-order transmission kinetics. PLoS ONE, 16(3):e0247512, March 2021.

[5] Jatin Narula, Abhinav Tiwari, and Oleg A. Igoshin. Role of Autoregulation and Relative Synthesis of Operon Partners in Alternative Sigma Factor Networks. PLoS Computational Biology, 12(12):e1005267, 2016.

[6] Ottar N. Bjørnstad, Katriona Shea, Martin Krzywinski, and Naomi Altman. The SEIRS model for infectious disease dynamics. Nature Methods, 17(6):557–558, June 2020. Number: 6 Publisher: Nature Publishing Group.

[7] Christian P Schwall, Torkel E Loman, Bruno M C Martins, Sandra Cortijo, Casandra Villava, Vassili Kusmartsev, Toby Livesey, Teresa Saez, and James C W Locke. Tunable phenotypic variability through an autoregulatory alternative sigma factor circuit. Molecular Systems Biology, 17(7):1–16, 2021.

[8] Daniel T. Gillespie. The chemical Langevin equation. Journal of Chemical Physics, 113(1):297–306, 2000.

[9] Daniel T. Gillespie. A General Method for Numerically Simulating the Stochastic Time Evolution of Coupled Chemical Reactions. Journal of Computational Physics, 22:403–434, 1976.

[10] Daniel T. Gillespie. Exact stochastic simulation of coupled chemical reactions. Journal of Physical Chemistry, 81(25):2340–2361, 1977.

[11] James Schaff, Charles C. Fink, Boris Slepchenko, John H. Carson, and Leslie M. Loew. A general computational framework for modeling cellular structure and function. Biophysical Journal, 73(3):1135–1146, 1997.

[12] Stefan Hoops, Ralph Gauges, Christine Lee, Jürgen Pahle, Natalia Simus, Mudita Singhal, Liang Xu, Pedro Mendes, and Ursula Kummer. COPASI - A COmplex PAthway SImulator. Bioinformatics, 22(24):3067–3074, 2006.

[13] A. Gonzalez Gonzalez, A. Naldi, L. Sánchez, D. Thieffry, and C. Chaouiya. GINsim: A software suite for the qualitative modelling, simulation and analysis of regulatory networks. BioSystems, 84(2):91–100, 2006.

[14] Steffen Klamt, Julio Saez-Rodriguez, and Ernst D Gilles. Structural and functional analysis of cellular networks with CellNetAnalyzer. New York, 13:1–13, 2008.

[15] Lucian P. Smith, Frank T. Bergmann, Deepak Chandran, and Herbert M. Sauro. Antimony: a modular model definition language. Bioinformatics, 25(18):2452–2454, September 2009.

[16] Camille Terfve, Thomas Cokelaer, David Henriques, Aidan MacNamara, Emanuel Goncalves, Melody K. Morris, Martijn van Iersel, Douglas A. Lauffenburger, and Julio Saez-Rodriguez. CellNOptR: A flexible toolkit to train protein signaling networks to data using multiple logic formalisms. BMC Systems Biology, 6, 2012.

[17] Carlos F. Lopez, Jeremy L. Muhlich, John A. Bachman, and Peter K. Sorger. Programming biological models in Python using PySB. Molecular Systems Biology, 9(1):1–19, 2013.

[18] Jorn Starruß, Walter De Back, Lutz Brusch, and Andreas Deutsch. Morpheus: A user-friendly modeling environment for multiscale and multicellular systems biology. Bioinformatics, 30(9):1331–1332, 2014.

[19] Brian Drawert, Andreas Hellander, Ben Bales, Debjani Banerjee, Giovanni Bellesia, Bernie J. Daigle Jr, Geoffrey Douglas, Mengyuan Gu, Anand Gupta, Stefan Hellander, Chris Horuk, Dibyendu Nath, Aviral Takkar, Sheng Wu, Per Lötstedt, Chandra Krintz, and Linda R. Petzold. Stochastic Simulation Service: Bridging the Gap between the Computational Expert and the Biologist. PLOS Computational Biology, 12(12):e1005220, December 2016. Publisher: Public Library of Science.

[20] Atefeh Kazeroonian, Fabian Fröhlich, Andreas Raue, Fabian J. Theis, and Jan Hasenauer. CERENA: ChEmical REaction Network Analyzer—a toolbox for the simulation and analysis of stochastic chemical kinetics. PLOS ONE, 11(1):1–15, 01 2016.

[21] Leonard A. Harris, Justin S. Hogg, José Juan Tapia, John A.P. Sekar, Sanjana Gupta, Ilya Korsunsky, Arshi Arora, Dipak Barua, Robert P. Sheehan, and James R. Faeder. BioNetGen 2.2: Advances in rule-based modeling. Bioinformatics, 32(21):3366–3368, 2016.

[22] Oleksandr Ostrenko, Pietro Incardona, Rajesh Ramaswamy, Lutz Brusch, and Ivo F. Sbalzarini. pSSAlib: The partial-propensity stochastic chemical network simulator. PLOS Computational Biology, 13(12):e1005865, December 2017. Publisher: Public Library of Science.

[23] J. Kyle Medley, Kiri Choi, Matthias König, Lucian Smith, Stanley Gu, Joseph Hellerstein, Stuart C. Sealfon, and Herbert M. Sauro. Tellurium notebooks-An environment for reproducible dynamical modeling in systems biology. PLOS Computational Biology, 14(6):e1006220, June 2018. Publisher: Public Library of Science.

[24] Kiri Choi, J. Kyle Medley, Matthias König, Kaylene Stocking, Lucian Smith, Stanley Gu, and Herbert M. Sauro. Tellurium: An extensible python-based modeling environment for systems and synthetic biology. Biosystems, 171:74–79, September 2018.

[25] Zachary B. Haiman, Daniel C. Zielinski, Yuko Koike, James T. Yurkovich, and Bernhard O. Palsson. MASSpy: Building, simulating, and visualizing dynamic biological models in Python using mass action kinetics. PLOS Computational Biology, 17(1):e1008208, January 2021. Publisher: Public Library of Science.

[26] William Poole, Ayush Pandey, Andrey Shur, Zoltan A. Tuza, and Richard M. Murray. BioCRNpyler: Compiling chemical reaction networks from biomolecular parts in diverse contexts. PLOS Computational Biology, 18(4):e1009987, April 2022. Publisher: Public Library of Science.

[27] Jeff Bezanson, Stefan Karpinski, Viral B. Shah, and Alan Edelman. Julia: A Fast Dynamic Language for Technical Computing. arXiv, pages 1–27, 2012.

[28] Jeff Bezanson, Alan Edelman, Stefan Karpinski, and Viral B. Shah. Julia: A fresh approach to numerical computing. SIAM Review, 59(1):65–98, 2017.

[29] Yingbo Ma, Shashi Gowda, Ranjan Anantharaman, Chris Laughman, Viral Shah, and Chris Rackauckas. Modeling-toolkit: A composable graph transformation system for equation-based modeling, 2021. arXiv:2103.05244.

[30] Shashi Gowda, Yingbo Ma, Alessandro Cheli, Maja Gwóźzdź, Viral B. Shah, Alan Edelman, and Christopher Rackauckas. High-performance symbolic-numerics via multiple dispatch. ACM Commun. Comput. Algebra, 55(3):92–96, jan 2022.

[31] Christopher Rackauckas and Qing Nie. DifferentialEquations.jl – A Performant and Feature-Rich Ecosystem for Solving Differential Equations in Julia. Journal of Open Research Software, 5(15):15, 2017.

[32] Ciaran Welsh, Jin Xu, Lucian Smith, Matthias König, Kiri Choi, and Herbert M Sauro. libRoadRunner 2.0: a high performance SBML simulation and analysis library. Bioinformatics, 39(1):btac770, January 2023.

[33] John H. Abel, Brian Drawert, Andreas Hellander, and Linda R. Petzold. GillesPy: A Python Package for Stochastic Model Building and Simulation. IEEE Life Sciences Letters, 2(3):35–38, 2017.

[34] Davan Harrison. A Brief Introduction to Automatic Differentiation for Machine Learning, October 2021. arXiv:2110.06209 [cs].

[35] N. Obatake, A. Shiu, X. Tang, and A. Torres. Oscillations and bistability in a model of ERK regulation. J Math Biol, 79(4):1515–1549, 09 2019.

[36] A. Jain and P. Lang. SBMLToolkit.jl. https://github.com/SciML/SBMLToolkit.jl, 2022.

[37] S. A. Isaacson. ReactionNetworkImporters.jl. https://github.com/SciML/ReactionNetworkImporters.jl, 2022.

[38] René Lefever, Grégoire Nicolis, and Pierre Borckmans. The brusselator: it does oscillate all the same. Journal of the Chemical Society, Faraday Transactions 1: Physical Chemistry in Condensed Phases, 84(4):1013, 1988.

[39] Yuri A. Kuznetsov. Elements of Applied Bifurcation Theory, volume 112 of Applied Mathematical Sciences. Springer, New York, NY, 2004.

[40] José M.G. Vilar, Hao Yuan Kueh, Naama Barkai, and Stanislas Leibler. Mechanisms of noise-resistance in genetic oscillators. Proceedings of the National Academy of Sciences of the United States of America, 99(9):5988–5992, 2002.

[41] Simon Christ, Daniel Schwabeneder, Christopher Rackauckas, Michael Krabbe Borregaard, and Thomas Breloff. Plots.jl – a user extendable plotting api for the julia programming language, 2022.

[42] Melanie I. Stefan, Thomas M. Bartol, Terrence J. Sejnowski, and Mary B. Kennedy. Multi-state modeling of biomolecules. PLOS Computational Biology, 10(9):1–9, 09 2014.

[43] Joshua Colvin, Michael I. Monine, James R. Faeder, William S. Hlavacek, Daniel D. Von Hoff, and Richard G. Posner. Simulation of large-scale rule-based models. Bioinformatics, 25(7):910–917, 02 2009.

[44] Michael L. Blinov, James R. Faeder, Byron Goldstein, and William S. Hlavacek. A network model of early events in epidermal growth factor receptor signaling that accounts for combinatorial complexity. Biosystems, 83(2):136–151, 2006. 5th International Conference on Systems Biology.

[45] Dipak Barua, William S. Hlavacek, and Tomasz Lipniacki. A computational model for early events in b cell antigen receptor signaling: Analysis of the roles of lyn and fyn. The Journal of Immunology, 189(2):646–658, 2012.

[46] James R. Faeder, William S. Hlavacek, Ilona Reischl, Michael L. Blinov, Henry Metzger, Antonio Redondo, Carla Wofsy, and Byron Goldstein. Investigation of early events in fcεri-mediated signaling using a detailed mathematical model. The Journal of Immunology, 170(7):3769–3781, 2003.

[47] Abhishekh Gupta and Pedro Mendes. An Overview of Network-Based and -Free Approaches for Stochastic Simulation of Biochemical Systems. Computation (Basel), 6(1), 2018.

[48] Christopher Rackauckas and Qing Nie. Adaptive methods for stochastic differential equations via natural embeddings and rejection sampling with memory. Discrete Continuous Dyn Syst Ser B., 22(7):2731–2761, 2017.

[49] Giulia Simoni, Federico Reali, Corrado Priami, and Luca Marchetti. Stochastic simulation algorithms for computational systems biology: Exact, approximate, and hybrid methods. Wiley Interdisciplinary Reviews: Systems Biology and Medicine, 11(6):e1459, 2019.

[50] S. A. Isaacson, V. Ilin, and C. V. Rackauckas. JumpProcesses.jl. https://github.com/SciML/JumpProcesses.jl/, 2022.

[51] J. M. McCollum, G. D. Peterson, C. D. Cox, M. L. Simpson, and N. F. Samatova. The sorting direct method for stochastic simulation of biochemical systems with varying reaction execution behavior. Computational Biology and Chemistry, 30(1), 2006.

[52] C. V. Rackauckas. Differentialequations.jl documentation. https://diffeq.sciml.ai/stable/.

[53] Ido Golding, Johan Paulsson, Scott M. Zawilski, and Edward C. Cox. Real-time kinetics of gene activity in individual bacteria. Cell, 123(6):1025–1036, 2005.

[54] Augustinas Sukys and Ramon Grima. MomentClosure.jl: automated moment closure approximations in Julia. Bioinformatics, 38(1):289–290, 06 2021.

[55] Kaan Öcal and Augustinas Sukys. FiniteStateProjection.jl. https://github.com/kaandocal/FiniteStateProjection.jl, 2022.

[56] Xiaoming Fu, Xinyi Zhou, Dongyang Gu, Zhixing Cao, and Ramon Grima. DelaySSAToolkit.jl: Stochastic simulation of reaction systems with time delays in Julia. Bioinformatics, 07 2022.

[57] Paul Breiding and Sascha Timme. HomotopyContinuation.jl: A Package for Homotopy Continuation in Julia. In International Congress on Mathematical Software, pages 458–465. Springer, 2018.

[58] Hongyu Miao, Xiaohua Xia, Alan S. Perelson, and Hulin Wu. On identifiability of nonlinear ode models and applications in viral dynamics. SIAM Review, 53(1):3–39, 2011.

[59] Hong Ge, Kai Xu, and Zoubin Ghahramani. Turing: a language for flexible probabilistic inference. In International Conference on Artificial Intelligence and Statistics, AISTATS 2018, 9-11 April 2018, Playa Blanca, Lanzarote, Canary Islands, Spain, pages 1682–1690, 2018.

[60] Christopher Rackauckas, Yingbo Ma, Julius Martensen, Collin Warner, Kirill Zubov, Rohit Supekar, Dominic Skinner, Ali Ramadhan, and Alan Edelman. Universal Differential Equations for Scientific Machine Learning, November 2021. arXiv:2001.04385 [cs, math, q-bio, stat].

[61] Romain Veltz. BifurcationKit.jl. https://hal.archives-ouvertes.fr/hal-02902346, mJul 2020. Package version: 0.1.8.

[62] Vaibhav Kumar Dixit and Christopher Rackauckas. Globalsensitivity.jl: Performant and parallel global sensitivity analysis with julia. Journal of Open Source Software, 7(76):4561, 2022.

[63] Benjamin Hepp, Ankit Gupta, and Mustafa Khammash. Adaptive hybrid simulations for multiscale stochastic reaction networks. The Journal of Chemical Physics, 142(3):034118, January 2015. Publisher: American Institute of Physics.

[64] Stefanie Winkelmann and Christof Schütte. Hybrid models for chemical reaction networks: Multiscale theory and application to gene regulatory systems. The Journal of Chemical Physics, 147(11):114115, September 2017. Publisher: American Institute of Physics.

[65] Daniel T. Gillespie. Approximate accelerated stochastic simulation of chemically reacting systems. The Journal of Chemical Physics, 115(4):1716–1733, July 2001.

[66] Yang Cao, Daniel T. Gillespie, and Linda R. Petzold. Avoiding negative populations in explicit Poisson tau-leaping. The Journal of Chemical Physics, 123(5):054104, August 2005.

[67] D. F. Anderson, D. J. Higham, S. C. Leite, and R. J. Williams. On constrained Langevin equations and (bio)chemical reaction networks. Multiscale Model. Simul., 17(1):1–30, 2018.

[68] D. J. Higham. Modeling and simulating chemical reactions. SIAM Review, 50(2):347–368, 2008.

[69] Jiahao Chen and Jarrett Revels. Robust benchmarking in noisy environments. arXiv e-prints, Aug 2016.

[70] Linda Petzold. Automatic selection of methods for solving stiff and nonstiff systems of ordinary differential equations. SIAM Journal on Scientific and Statistical Computing, 4(1):136–148, 1983.

[71] Alan C Hindmarsh, Peter N Brown, Keith E Grant, Steven L Lee, Radu Serban, Dan E Shumaker, and Carol S Wood-ward. SUNDIALS: Suite of nonlinear and differential/algebraic equation solvers. ACM Transactions on Mathematical Software (TOMS), 31(3):363–396, 2005.

[72] ME Hosea and LF Shampine. Analysis and implementation of tr-bdf2. Applied Numerical Mathematics, 20(1-2):21–37, 1996.

[73] Lawrence F Shampine and Mark W Reichelt. The matlab ode suite. SIAM journal on scientific computing, 18(1):1–22, 1997.

[74] Albert Reuther, Jeremy Kepner, Chansup Byun, Siddharth Samsi, William Arcand, David Bestor, Bill Bergeron, Vijay Gadepally, Michael Houle, Matthew Hubbell, Michael Jones, Anna Klein, Lauren Milechin, Julia Mullen, Andrew Prout, Antonio Rosa, Charles Yee, and Peter Michaleas. Interactive supercomputing on 40,000 cores for machine learning and data analysis. In 2018 IEEE High Performance extreme Computing Conference (HPEC), pages 1–6. IEEE, 2018.

[75] Vo Hong Thanh, Corrado Priami, and Roberto Zunino. Efficient rejection-based simulation of biochemical reactions with stochastic noise and delays. The Journal of Chemical Physics, 141(13):134116–134113, 2014.

[76] Vo Hong Thanh, Roberto Zunino, and Corrado Priami. On the rejection-based algorithm for simulation and analysis of large-scale reaction networks. The Journal of Chemical Physics, 142(24):244106–244114, 2015.

[77] Vo Hong Thanh, Roberto Zunino, and Corrado Priami. Efficient constant-time complexity algorithm for stochastic simulation of large reaction networks. IEEE/ACM Transactions on Computational Biology and Bioinformatics, 14(3):657–667, 2017.

